# A kinetic ruler controls mRNA poly(A) tail length

**DOI:** 10.1101/2025.05.01.651612

**Authors:** Emilie Gabs, Emil Aalto-Setälä, Aada Välisaari, Anssi M. Malinen, Torben Heick Jensen, Stephen H. McLaughlin, Lori A. Passmore, Matti Turtola

**Affiliations:** Department of Life Technologies, University of Turku, Finland; Department of Molecular Biology and Genetics, Aarhus University, Denmark; Laboratory of Molecular Biology, Cambridge, United Kingdom

## Abstract

Poly(A) tails of newly synthesized mRNAs have uniform lengths, arising through cooperation between the cleavage and polyadenylation complex (CPAC) and poly(A) binding proteins (PABPs). In the budding yeast, *Saccharomyces cerevisiae*, the responsible PABP is the evolutionarily conserved CCCH zinc finger protein Nab2 that facilitates the biogenesis of ∼60 adenosine mRNA poly(A) tails. Here, we address the molecular basis for such length control. Reconstituting polyadenylation reactions during the formation of Nab2:poly(A) RNA ribonucleoprotein particles *in vitro*, we find that Nab2 dimerization directs polyadenylation termination. The Nab2 dimer is stable only on poly(A) tails that are longer than 25 adenosines, explaining how Nab2 avoids prematurely terminating poly(A) synthesis. However, the mature tail length is not determined by the footprint of Nab2 on the RNA, but rather by the kinetic competition between CPAC-mediated tail elongation and Nab2 RNA-binding. Variations in Nab2 RNA-binding rate can shift poly(A) tail lengths, but in cells such variations are buffered by autoregulation of Nab2 protein concentration. As a result, poly(A) tail length control operates through a “kinetic ruler” mechanism, whereby the concentration of Nab2 quantifies RNA length.

## INTRODUCTION

RNA 3′end polyadenosine (poly(A)) tails affect all main post-transcriptional steps of gene expression. A significant property of the poly(A) tail is its length, which impacts the recruitment of poly(A) binding proteins (PABPs) and which is functionally linked to export, translatability and stability of the RNA (Passmore and Coller 2021). Poly(A) tails of mRNAs are synthesized by the multisubunit cleavage and polyadenylation complex (CPAC), and because the reaction is non-templated, the product length must be controlled by auxiliary factors, which are largely PABPs (reviewed in (Boreikaite and Passmore 2023; Eckmann et al. 2011; Rodríguez-Molina and Turtola 2023)). As a result, all polyadenylated mRNAs are initially produced with a species-characteristic tail length, with newly synthesized yeast and mammalian poly(A) tails having ∼60 and ∼250 adenosines, respectively.

From its uniform starting point, the length of the poly(A) tail of a given mRNA varies throughout its lifecycle as a result of nuclear and cytoplasmic activities that either extend or shorten the tail (Passmore and Coller 2021; Eckmann et al. 2011; Xiang et al. 2024; Lim et al. 2016; Tudek et al. 2021; Eisen et al. 2020; Alles et al. 2023). Thus, the steady-state poly(A) tail profile reflects its different cellular phases. Poly(A) tail length control provides a uniform entry point for deadenylases that gradually shorten the poly(A) tail with variable, transcript- and condition-specific rates (Passmore and Coller 2021). Deadenylation generally stimulates full mRNA degradation (Czarnocka-Cieciura et al. 2024; Parker 2012; Webster et al. 2018; Eisen et al. 2020), often by promoting decapping, which can also be a rate-limiting step (Audebert et al. 2024; Wiener et al. 2021). Initial poly(A) tail length is therefore a key parameter in determining mRNA abundance.

In *S. cerevisiae*, the nuclear PABP Nab2 is the primary factor controlling mRNA poly(A) tail length. It does so by associating with the growing poly(A) tail and inhibiting the polyadenylation activity of the CPAC (Hector et al. 2002; Turtola et al. 2021; Viphakone et al. 2008). Nab2 binding is further coupled to the formation of an export-competent messenger ribonucleoprotein particle (mRNP), as Nab2 interacts with proteins required for mRNP export (Iglesias et al. 2010; Bonneau et al. 2023; Carmody et al. 2010; Asada et al. 2023), facilitates mRNP docking to the nuclear pore complex (Saroufim et al. 2015; Fasken et al. 2008) and, during its residence in the nucleus, protects the newly synthesized transcript from degradation (Schmid et al. 2015; Tudek et al. 2018; Turtola et al. 2021). Conversely, errors in mRNP assembly or export disrupt Nab2-mediated poly(A) tail length control and lead to rapid degradation of mis-processed transcripts (Hilleren and Parker 2001; Jensen et al. 2001; Libri et al. 2002; Rougemaille et al. 2007; Tudek et al. 2018; Turtola et al. 2021). Consequently, Nab2 function is required for the proper production of RNA polymerase II (Pol II) transcribed mRNAs (González-Aguilera et al. 2011; Schmid et al. 2015).

The Nab2 homologous protein in animals, ZC3H14, has been reported to associate with mRNP biogenesis factors and components of the spliceosome (Morris and Corbett 2018; Soucek et al. 2016; Li et al. 2024), to impact alternative splicing (Jalloh et al. 2023) and to promote circular RNA generation through back-splicing (Li et al. 2024). Moreover, ZC3H14 has been implicated in the surveillance of non-coding RNAs (Latour et al. 2025), and the turnover of prematurely terminated Pol II transcripts (Insco et al. 2023). However, depletion studies have also pointed to ZC3H14 restricting poly(A) tail length (Pak et al. 2011; Kelly et al. 2014; Rha et al. 2017; Bienkowski et al. 2017; Morris and Corbett 2018; Li et al. 2024). Loss of the ZC3H14 homologue in *Drosophila*, dNab2, decreases viability and causes defective neuronal development (Kelly et al. 2014, 2016; Pak et al. 2011). Likewise, in mouse models, loss of functional *Zc3h14* impairs neuronal development (Rha et al. 2017) and disrupts spermatogenesis (Li et al. 2024), whereas loss-of-function mutations in human *ZC3H14* lead to intellectual disability (Pak et al. 2011). Finally, in *C. elegans* and mouse brains the protein affects the susceptibility to pathological tau, hence, its alternative name ‘suppressor of tau 2 (SUT-2/MSUT2)’ (Guthrie et al. 2009; Wheeler et al. 2019). Altogether, these diverse phenotypes await mechanistic explanations as to how Nab2/ZC3H14 regulates different RNA processing activities. This might be more easily revealed in organisms such as *S. cerevisiae*, in which a limited set of nuclear PABPs simplifies the analysis of poly(A)-tail mediated regulation.

PABP-mediated regulation of enzymes that either extend or shorten poly(A) tails is tied to their length-dependent interaction with poly(A) RNA. The current picture of how this is achieved comes from studies on RNA recognition motif (RRM)-containing PABPs, including mammalian PABPN1 and PABPC. These PABPs align their RRM domains side-by side along the poly(A) RNA chain, like beads on a string (Eckmann et al. 2011; Passmore and Coller 2021). Consequently, the RNA footprints of the RRMs (11 adenosines for PABPN1 (Meyer et al. 2002) and 27 adenosines for PABPC (Baer and Kornberg 1980)) translate into periodic *molecular rulers* that control enzyme processivities in poly(A) tail length-dependent manners. In the case of PABPN1, the assembly of 15-20 PABPN1 molecules on a ∼250 adenosine long poly(A) tail collapses the RNP into a globular structure, which is suggested to terminate processive polyadenylation by disrupting the interactions between the poly(A) tail-bound PABPN1, the poly(A) polymerase (PAP) and the rest of the CPAC machinery (Kühn et al. 2009, 2017). On the other hand, the cooperation of the poly(A)-bound PABPC and exonuclease activities results in poly(A) tail shortening to lengths that have ∼30 adenosine periodicity (Webster et al. 2018; Schäfer et al. 2019; Azoubel Lima et al. 2017). However, similar structure-function relationship has not yet been established for structurally different Nab2/ZC3H14 PABPs, which belong to Cys-Cys-Cys-His (CCCH) type zinc finger (ZnF) proteins.

Compared to RRM-type PABPs, the CCCH ZnF Nab2 has a profoundly different topology for interacting with poly(A) RNA. It displays a diffuse nuclease-protected poly(A) RNA footprint (Viphakone et al. 2008), it harbours structurally non-similar tandem ZnF domains that mediate RNA binding (Brockmann et al. 2012; Martínez-Lumbreras et al. 2013), and it employs an unusual mode of RNA interaction where a dimeric interface between three ZnFs from two Nab2 molecules creates a binding surface for 7 adenosines (Aibara et al. 2017). The functional significance of this dimeric architecture remains elusive and is therefore at present insufficient to explain how Nab2/ZC3H14 proteins are able to specify mRNA poly(A) tail lengths.

Here, we examine the biochemical basis of how Nab2, the founding member of CCCH ZnF PABPs (Fasken et al. 2019), interacts with poly(A) RNA to regulate its tail length during CPAC-mediated polyadenylation reaction. We find that, while Nab2 dimerization and its multidomain RNA-binding mode sensitize its interaction to different poly(A) RNA lengths, its footprint on poly(A) RNA does not readily determine poly(A) tail lengths. Rather, such control relies on the fine-tuned kinetics of Nab2:poly(A) binding that competes with the poly(A) tail synthesis by the CPAC. We propose that Nab2 acts as a concentration-dependent *kinetic ruler* that quantifies RNA chain length.

## RESULTS

### Time-resolved measurements of poly(A) tail elongation by the CPAC reveal distinct reaction phases

To obtain a detailed understanding of Nab2-mediated poly(A) tail length control, we first characterized the kinetics of CPAC-mediated polyadenylation reactions. To this end, polyadenylation-competent CPAC was reconstituted using purified CPF, CF IA and CF IB subcomplexes (**Fig. 1A**). As a polyadenylation substrate, we used 42-nucleotide RNA fragment from the *CYC1* 3′UTR (*CYC1*_*42*_) labelled with an Atto680 fluorescent dye at the 5′end. *CYC1*_*42*_ contained the polyadenylation signal sequence (PAS) that is recognized by CPF and the flanking sequences that are bound by CF IA and CF IB (Hill et al. 2019; Dichtl and Keller 2001). The 3′end of *CYC1*_*42*_ constituted the CPF-mediated endonucleolytic cleavage site, and therefore this RNA could be used as a ‘pre-cleaved’ substrate for the isolated polyadenylation reaction (Schmid et al. 2012; Viphakone et al. 2008).

**Figure 1.**
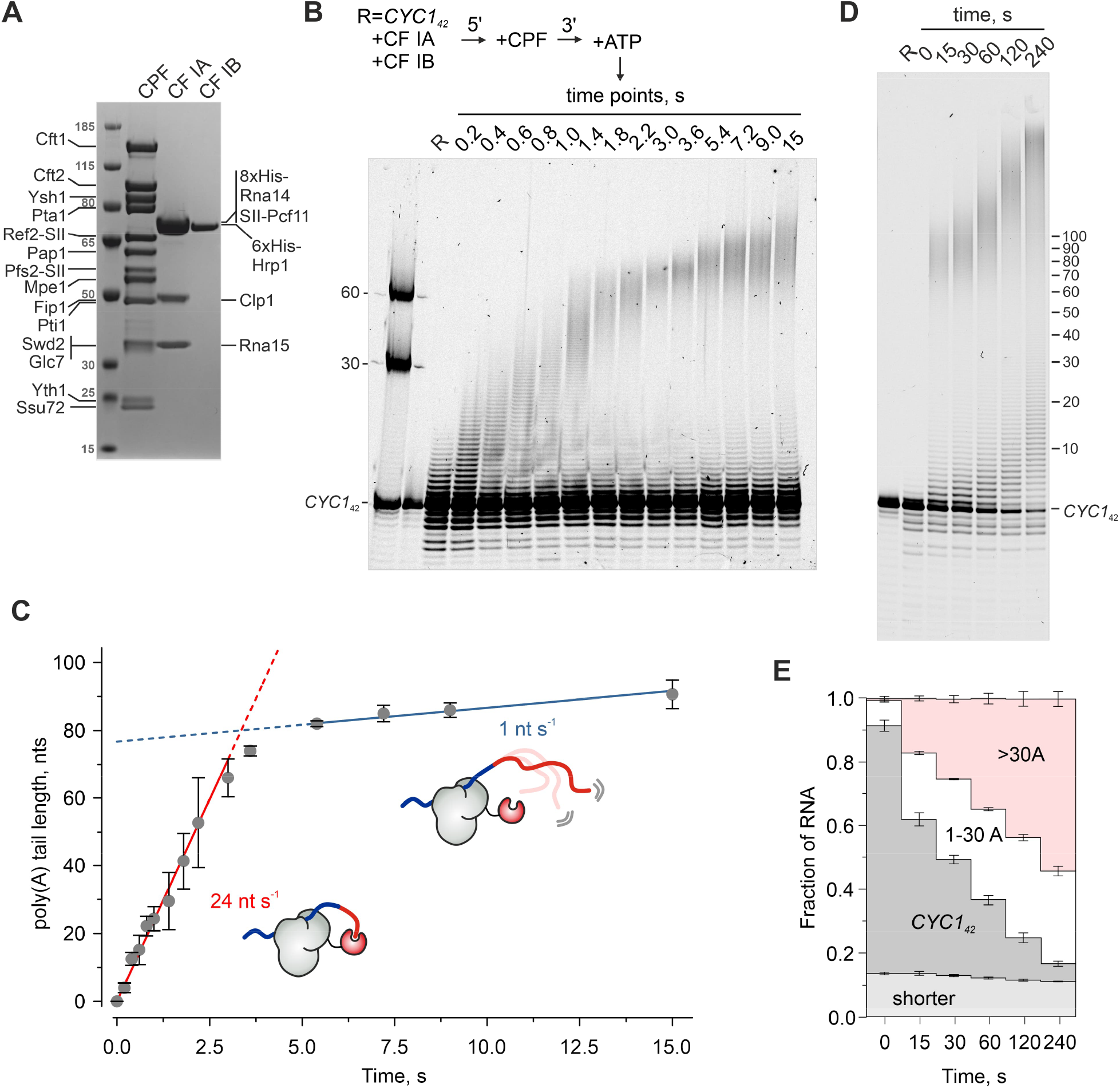
Time-resolved measurement of RNA polyadenylation by the CPAC. (*A*) SDS-PAGE of CPAC proteins used for reconstituting the polyadenylation reactions. (*B*) Time-resolved poly(A) tail elongation of the *CYC1*_*42*_ RNA by the CPAC. Reactions were initiated by mixing the pre-assembled CPAC with ATP (2 mM) in the quench flow apparatus and stopped with HCl at indicated times. Final concentrations were: *CYC1*_*42*_ (50 nM), CPF (50 nM), CF IA (225 nM) and CF IB (225 nM). The 5′Atto680-labelled RNA reaction products were resolved on a denaturing polyacrylamide gel and detected by a fluorescent scanner. The size markers indicate the lengths of poly(A) tails. (*C*) Quantification of poly(A) tail elongation rate. Mean poly(A) tail lengths (see Methods) averaged from three time-resolved measurements are displayed with standard deviations. The rates of poly(A) tail elongation were calculated from the slope of the linear fit to the first ten time points for the initial 70 adenosines (red continuous line), and from the last four time points for the elongation after the addition of 80 adenosines (blue continuous line). The schematic drawings present an interpretation for the fast and slow modes of poly(A) tail elongation by the CPAC (grey; poly(A) polymerase in red). (*D*) Longer time course of *CYC1*_*42*_ RNA polyadenylation with the same reaction conditions as in *B*. (*E*) Quantification of polyadenylation reaction products at the time points in *D*, showing fractional amounts of RNA species (dark grey, non-processed *CYC1*_*42*_; white, 1-30 adenosine poly(A) tail; red, > 30 adenosine poly(A) tail; light grey, cleavage products) as stacked bars with standard deviations from three measurements.

Polyadenylation was initiated by the addition of ATP and samples were quenched at different time points to monitor reaction progression. In order to resolve poly(A) tail elongation kinetics, we used the quench flow technique for initiating and stopping reactions in a sub-second time scale. As shown in **Fig. 1B**, and quantified in **Fig. 1C**, a fraction of *CYC1*_*42*_ RNA was elongated by ∼70 adenosines with a rate of 24 nt s^-1^. These adenosines were added in a single wave and in an apparently processive manner, indicating a continuous engagement of the CPF and the poly(A) polymerase Pap1 throughout this phase of the reaction. We note because CF IA and CF IB are required for efficient polyadenylation (Turtola et al. 2021; Gross and Moore 2001; Casañal et al. 2017), the whole CPAC was evidently assembled on the RNA substrate. Notably, the rate of poly(A) tail elongation subsequently slowed down significantly, with tails >80 adenosines being elongated with a rate of 1 nt s^-1^. Tracking the reaction progress on a longer time scale showed that the slow extension of poly(A) tails continued far beyond 100 adenosines (**Fig. 1D**). We speculate that this slow mode of poly(A) tail elongation was due to the increased distance between the RNA 3′end and the CPF-bound Pap1, which is tethered to the PAS (see schematics in Fig. 1C). Here the increasing length of the poly(A) tail reduces the encounters between the RNA 3′end and the active site of Pap1.

The long time point experiment also revealed a group of short-tailed reaction products that were extended at an even slower rate, with poly(A) tails reaching ∼30 adenosines in 4 minutes, equalling an elongation rate of ∼0.1 nt s^-1^ (**Fig. 1D**). The proportion of these reaction products increased when there was an excess of *CYC1*_*42*_ over CPF, and they were elongated faster at high CPF concentrations (**Supplemental Fig. S1A**). This suggested distributive polyadenylation by CPF molecules that were not PAS-bound in *cis*, and appeared to be an artefact of using pre-cleaved RNA substrate, as it was not observed in coupled cleavage and polyadenylation reactions (**Supplemental Fig. S1B**).

Only ∼20% of the *CYC1*_*42*_ RNA was processively polyadenylated within the first 15 seconds, and it took several minutes for the reaction to consume all RNA substrate (**Fig. 1D-E**). This indicated a rate-limiting step before processive elongation. One possible rate-limiting step is the assembly of an elongation-competent CPAC. However, the resistance of a fraction of the *CYC1*_*42*_ substrate to both processive and distributive polyadenylation and its delayed polyadenylation at a later time point (e.g. compare lanes 4 and 5 in **Supplemental Fig. S1A**) hinted to an internal activation step within the CPAC. Notably, the start of polyadenylation appeared also slow in the coupled cleavage and polyadenylation reactions, as shown by the accumulation of 5′-cleavage products prior to their subsequent polyadenylation (**Supplemental Fig. S1B-C**).

In conclusion, CPAC-mediated polyadenylation *in vitro* can be described by three kinetic reaction phases: *i*) activation of polyadenylation, which limits the overall RNA processing rate, *ii*) rapid elongation at a rate of 24 nt s^-1^ until the poly(A) tail reaches ∼70 adenosines, *iii*) slowed elongation at a rate of ∼1 nt s^-1^ once tails are >80 adenosines. We note that the slow elongation of longer tails aligns with the previously reported “intrinsic” poly(A) tail length control, whereby the CPAC can restrict excessive polyadenylation in the absence of PABP regulation (Turtola et al., 2021).

### Poly(A) tail lengths are dictated by a kinetic competition between CPAC-mediated poly(A) tail elongation and Nab2 binding

To investigate how CPAC-mediated poly(A) tail synthesis is operated in the presence of a PABP, purified Nab2 (**Supplemental Fig. S2**) was included in the reactions. In the presence of 500 nM of Nab2, poly(A) tails were restricted to 50-70 adenosines all through the 4 min incubation time (**Fig. 2A**). Interestingly, when reactions were performed in the presence of varying amounts of Nab2, poly(A) tail lengths became progressively shorter with increasing Nab2 concentration (**Fig. 2B**). This implied that Nab2 binding kinetics play a role in length-determination.

**Figure 2.**
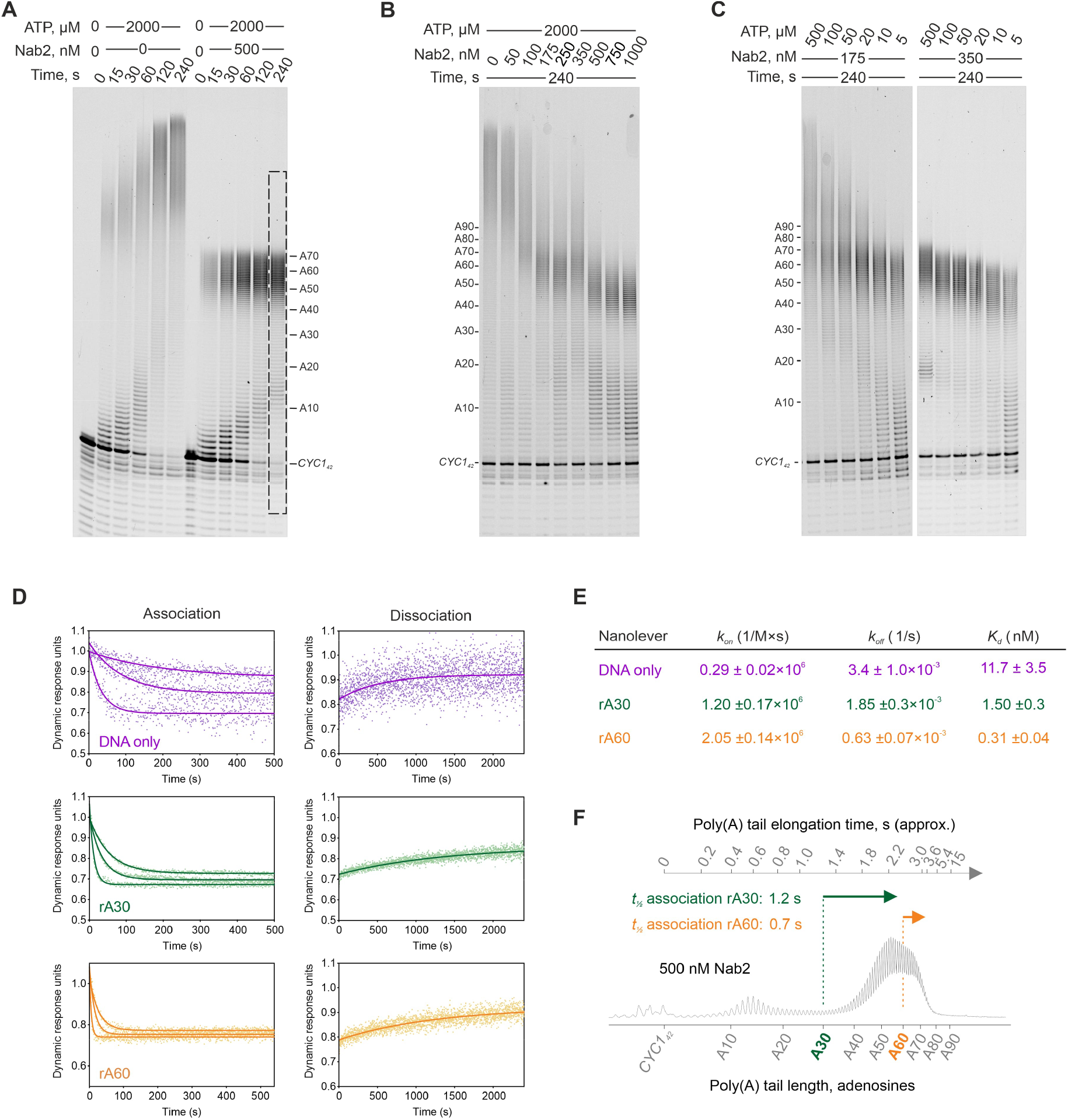
Poly(A) tail lengths are determined by a kinetic competition between CPAC-mediated poly(A) tail elongation and Nab2-mediated termination of poly(A) tail synthesis. (*A-C*) The effect of Nab2 on CPAC-mediated *CYC1*_*42*_ polyadenylation analysed at different time points (*A*), with varying concentrations of Nab2 (*B*), and with varying concentrations of ATP (*C*). The concentrations of *CYC1*_*42*_ (25 nM), CPF (75 nM), CF IA (500 nM) and CF IB (500 nM) were the same in panels *A-C*. The gel lane intensity scan of the marked area at 500 nM Nab2 is shown in *F*. (*D-E*) SwitchSENSE measurement of binding kinetics of Nab2 association with the poly(A) RNAs attached to the DNA nanolevers. Flowing buffer containing Nab2 at concentrations of 100, 33.3, 11.1 nM over the nanolevers attached to the chip surface decreases their switching speed when a complex is formed and is used to determine the association rate (*k*_*on*_). Dissociation of the complex is observed by flowing buffer over the chip surface resulting in an increase in the DNA switching speed and is used to determine the dissociation rate (*k*_*off*_). The dynamic response of the nanolevers versus time were used to fit association and dissociation curves of Nab2 with DNA nanolever only (magenta), A_30_ RNA (rA30, green) and A_60_ RNA (rA60, orange) to calculate average *k*_*on*_, *k*_*off*_ and *K*_*d*_ values and their corresponding standard errors shown in the table in *E*. (*F*) The poly(A) tail length profile from the highlighted gel area in *A* plotted along the approximated average poly(A) tail elongation time (from Fig. 1) and Nab2 association half-times (*t*_½_) to rA30 (green arrow) and rA60 (orange arrow) tails at 500 nM concentration of Nab2.

To test if Nab2-mediated poly(A) tail termination was sensitive to tail elongation rate, we performed polyadenylation reactions at different ATP concentrations. Notably, whereas 175 nM of Nab2 was unable to restrict polyadenylation at 500 µM of ATP, it effectively terminated poly(A) tail synthesis at lower ATP concentrations (**Fig. 2C**, left panel). Likewise, at 350 nM of Nab2, the modal poly(A) tail length decreased from ∼60 to ∼45 adenosines when the ATP concentration was decreased from 500 µM to 5 µM (**Fig. 2C**, right panel). Since the same concentration of Nab2 could give rise to different poly(A) tail lengths, this excluded control mechanisms that determine the poly(A) chain length by counting the number of nucleotides. Instead, this suggested that tail length control is kinetically determined as a result of the competition between the rates of poly(A) tail elongation and Nab2 RNA-binding.

To evaluate if Nab2 association dynamics were compatible with such a model, we used the SwitchSENSE technology to determine the association and dissociation rates of Nab2 binding to poly(A) RNAs that were linked to DNA nanolevers. The rate of Nab2 binding to a rA_60_ RNA (*k*_*on*_=2.05 ± 0.14 × 10^6^ M^-1^s^-1^) was nearly two times faster than to a rA_30_ RNA (*k*_*on*_=1.2 ± 0.17 × 10^6^ M^-1^s^-1^) and nearly ten times faster than to the DNA nanolever only (*k*_*on*_= 0.29 × 10^6^ M^-1^s^-1^) (**Fig. 2D-E**). Therefore, at a concentration of 500 nM of Nab2 the half-times (*t*_*1/2*_) of Nab2 association to rA_30_ and rA_60_ RNAs are ∼1.2 and ∼0.7 seconds, respectively. This is in good agreement with the duration of poly(A) tail elongation to 50-70 adenosines, which it takes the CPAC between 2-3 seconds to achieve (see Fig. 1C), and which are produced in the presence of 500 nM Nab2 (**Fig. 2F**).

We conclude that Nab2 does not operate as a molecular ruler to measure poly(A) tail length. Rather, the rates of poly(A) tail elongation and Nab2 RNA-binding govern a zone of termination, resulting in a restricted distribution of tail lengths. The slowdown of tail elongation after its first ∼70 adenosines efficiently reduces the likelihood of producing hyperadenylated poly(A) tails.

### Poly(A) tail length-dependent dimerization of Nab2

The dissociation constants (*K*_*d*_) derived from the Nab2 on- and off-rates indicated that the protein binds with ∼5-fold higher affinity to A_60_ RNA than to A_30_ RNA (**Fig. 2E**). This prompted us to study the biochemical properties of Nab2:poly(A) RNPs, which we first did using electromobility shift assays (EMSAs) of Nab2 and a fluorescently labelled A_59_ RNA. Notably, the gel shift pattern indicated that one, two or more copies of Nab2 bind to A_59_ RNA (**Fig. 3A**, *left*). The positive Hill coefficient (*n*_H_ = 2.7) suggested that cooperative interactions between monomeric units of Nab2 enhance its RNA-binding (**Fig. 3A**, *right*).

**Figure 3.**
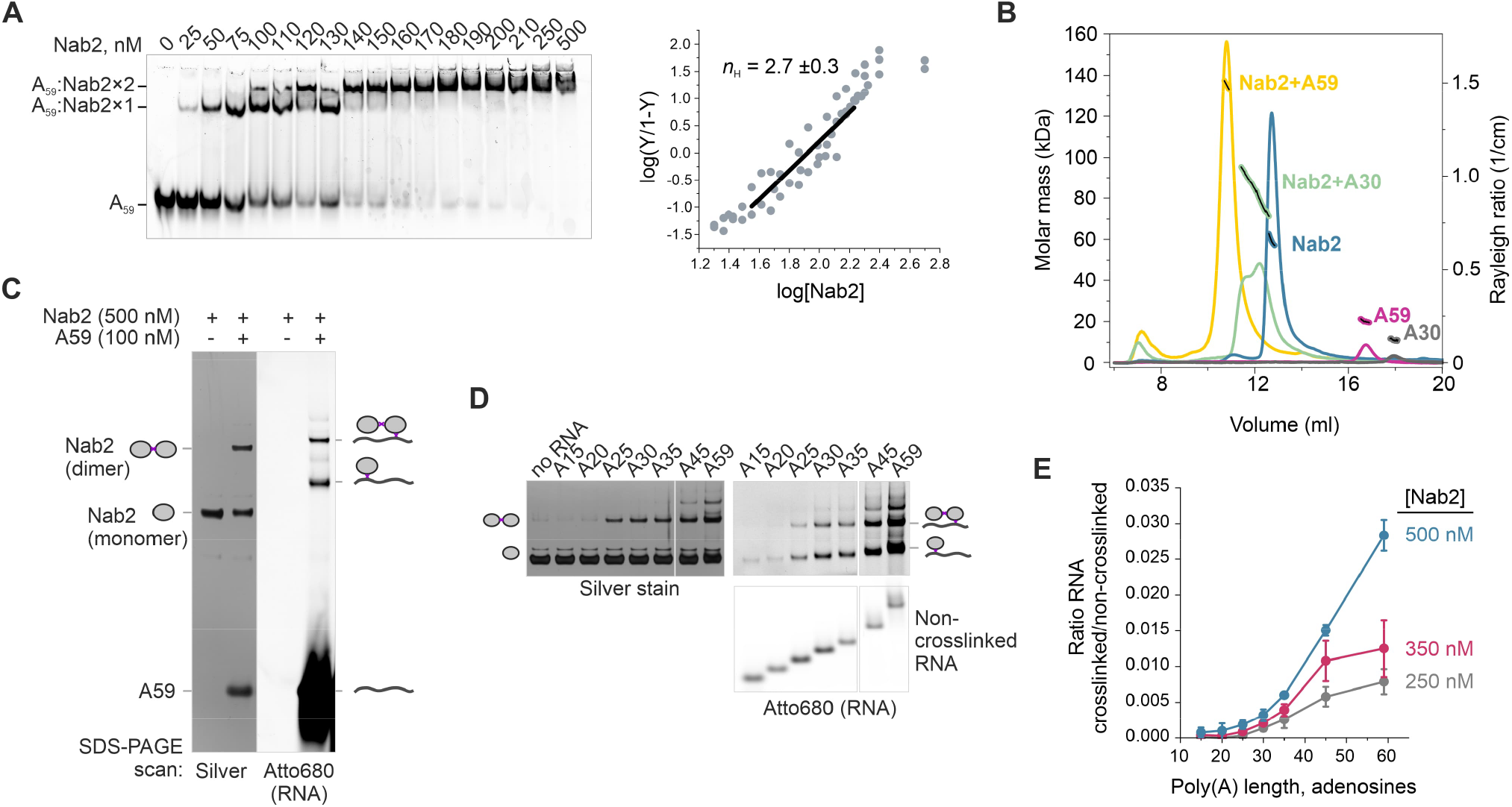
Poly(A) tail length-dependent dimerization of Nab2. (*A*) *Left:* Electrophoretic mobility shift assay for Nab2 interaction with Atto680-labelled A_59_ RNA. *Right:* The fractional saturation (Y) of Nab2 binding to A_59_ was quantified from three experiments and fitted to the Hill equation (see Methods) to calculate the Hill coefficient (*n*_H_ ± standard error). (*B*) SEC-MALS analysis of A_59_ (magenta), A_30_ (light grey), Nab2 (blue) and Nab2 mixed with either A_59_ (yellow) or A_30_ (green). Light scattering intensities are displayed (y-axis on the right side) along the elution volume (x-axis). The discontinuous black lines with coloured outlines show the calculated molecular masses, determined by the light scattering properties and analysed across the main peaks (y-axis on the left side). The calculated molecular weights are shown in **Supplemental Table S1**. (*C*) Formaldehyde crosslinking of Nab2 and Atto680-A_59_. Samples treated with 0.3% formaldehyde were quenched with glycine and separated by denaturing SDS-PAGE. The gels were scanned for Atto680 signal (right panel) and then stained with silver (left panel). (*D*) Formaldehyde crosslinking of Nab2 (500 nM) with Atto680-labelled poly(A) RNAs (100 nM) of varying lengths. The gel panels were cropped from the same gel. (*E*) Quantification of poly(A) RNA-Nab2 crosslinks at 500, 350 or 250 nM Nab2. The corresponding Atto680 scans and silver stained gel panels are shown in **Supplemental Fig. S3B**.

To determine the Nab2 oligomerization state more precisely, we used size exclusion chromatography coupled to multiangle light scattering (SEC-MALS). Nab2 (M_r_ 59.1 kDa) mixed with A_59_ (M_r_ 19.4 kDa) eluted as a complex with a molecular weight of 135 kDa, consistent with a complex stoichiometry of 2:1 Nab2:RNA, which was further supported by the conjugate analysis that separated the total molecular weight to protein and RNA components (**Fig. 3B, Supplemental Table S1**). Doubling the ratio of protein to RNA in the injected sample did not yield higher order complexes. Moreover, Nab2 or A_59_ injected alone eluted as monomeric species, demonstrating that Nab2 dimerization is dependent on poly(A) RNA. Interestingly, a complex formed between Nab2 and the A_30_ RNA (M_r_ 9.9 kDa) migrated as a bimodal peak between the dimeric and monomeric Nab2 species. Light scattering and UV-signals (**Supplemental Fig. S3A**) were consistent with one Nab2 binding either one, two or three molecules of A_30_ RNA (**Supplemental Table S1**), suggesting that the stability of the Nab2 dimer is sensitive to the length of poly(A) RNA.

In order to quantify such length dependence, we used formaldehyde crosslinking to capture the Nab2-poly(A) RNA interaction. As shown by silver staining in **Fig. 3C**, intermolecular crosslinks between two molecules of Nab2 were observed in the presence of the A_59_ RNA, but not in its absence. Crosslinks between Nab2 and A_59_ were also detected, although they formed with lower efficiencies. Because the efficiency of RNA-protein crosslinking was sensitive to the concentration of Nab2 (**Supplemental Fig. S3B**), we used it as a semi-quantitative measure of the interaction between Nab2 and poly(A) RNAs of varying lengths. Whereas 3% of the A_59_ RNA crosslinked to Nab2 (at 500 nM) within the 10 min incubation period, crosslinking was only 0.08% and 0.1% for the A_15_ and A_20_ RNAs, respectively (**Fig. 3D-E, Supplemental Fig. S3B**). Only after increasing poly(A) tail length to >25 adenosines, crosslinking efficiency increased and the formation of crosslinked Nab2-dimers followed the same trend. Thus, low crosslinking efficiency of short RNA could not be explained solely by less available crosslinking sites. These results therefore suggested that the formation of a Nab2-dimer requires a continuous poly(A) RNA, which is at least 25 adenosines long, and that the interaction between Nab2 and RNA becomes progressively more stable as the poly(A) tail gets longer.

### Dimerized Nab2 terminates polyadenylation

To investigate the functional significance of Nab2 dimerization, we used the rapamycin-dependent heterodimerization of the FKBP12 and FRB domains (Choi et al. 1996) to conditionally tether two Nab2 molecules together (**Fig. 4A**). We constructed and purified fusion proteins, where either FKBP12 or FRB domains were C-terminally linked to Nab2 (**Supplemental Fig. S2**). Mass photometry was used to verify that Nab2-FKBP12 and Nab2-FRB proteins were monomeric in solution but formed a heterodimeric complex in the presence of rapamycin (**Fig. 4B, Supplemental Fig. S4A**). In contrast, poly(A) RNA was required to link two unmodified Nab2 molecules within a ternary complex (**Supplemental Fig. S4A**).

**Figure 4.**
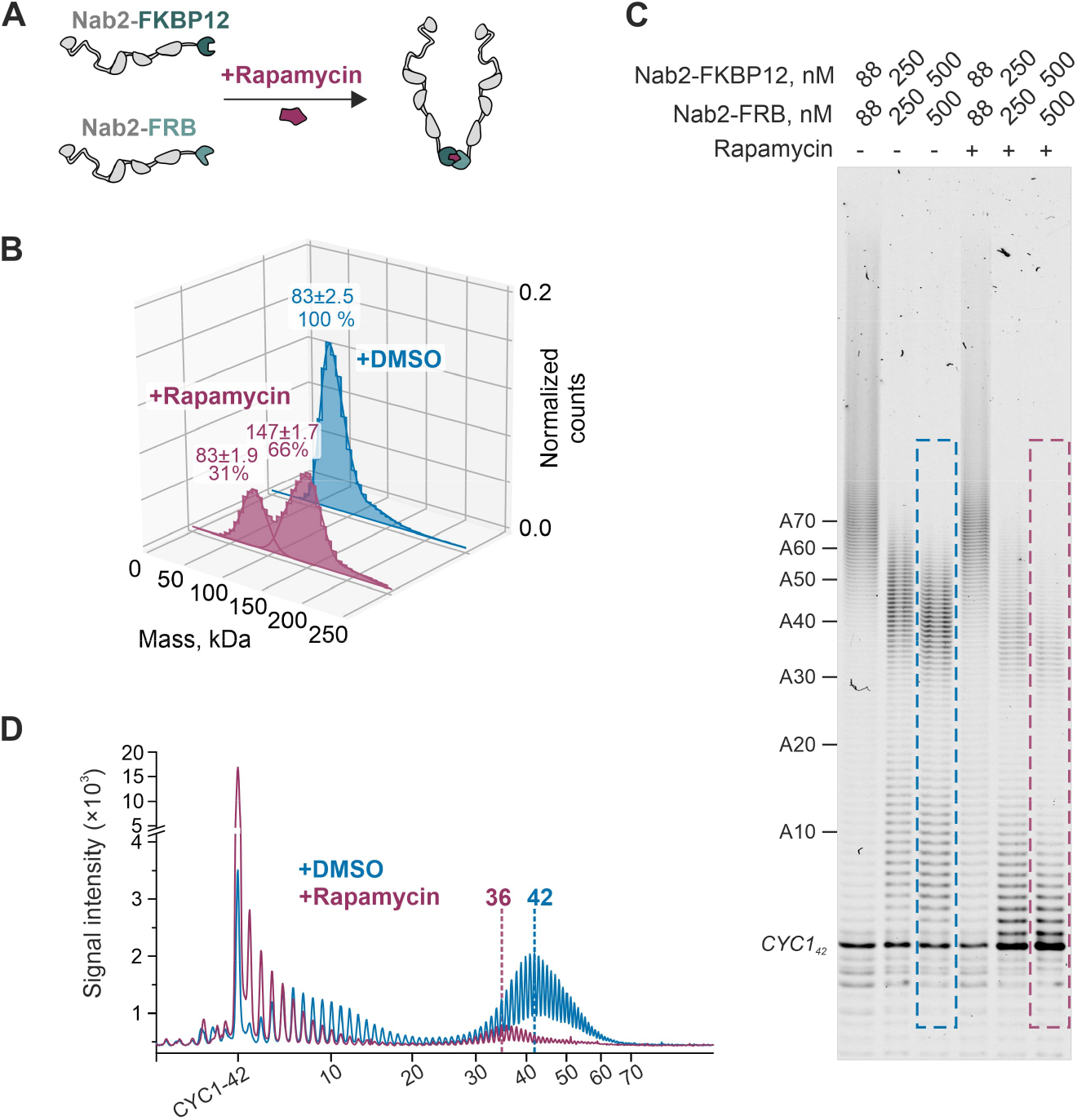
Dimerized Nab2 inhibits polyadenylation. (*A*) The principle of rapamycin induced dimerization of Nab2-FKBP12 and Nab2-FRB fusion proteins. (*B*) Mass photometric analysis of Nab2-FKBP12 (125 nM) and Nab2-FRB (125 nM) in the presence of DMSO (2.5 % v/v) or rapamycin (1 µM). The Gaussian fits of the count-normalized histograms are shown as lines with the peak position ± SEM (in kDa) and the percentage of total peak counts displayed above. (*C*) The effect of rapamycin-induced Nab2 dimerization on *CYC1*_*42*_ polyadenylation by the CPAC. Nab2-FKBP12 and Nab2-FRB were incubated with DMSO (2.5 % v/v; -rapamycin) or with 1 µM rapamycin before being added to the polyadenylation reactions together with ATP. The reactions were stopped after 4 minutes. (*D*) The gel lane intensity scans from the marked areas in *C*. The modal poly(A) tail lengths (beyond >30As) are marked with vertical lines. See **Supplemental Fig. S4** for additional controls.

Nab2-FKBP12 and Nab2-FRB proteins were capable of terminating CPAC-mediated polyadenylation, both independently and in combination (**Supplemental Fig. S4B**). Thus, the fused domains did not interfere with termination activity. Notably, when the Nab2-FKBP12 and Nab2-FRB proteins were linked together by rapamycin before the start of polyadenylation reactions, *CYC1*_*42*_ polyadenylation was largely prevented (**Fig. 4C**). Therefore, facilitated dimerization of Nab2 effectively inhibits polyadenylation of a non-polyadenylated substrate. Interestingly, a small fraction of substrate RNA that escaped the initial inhibition was polyadenylated to a modal length of ∼36 adenosines at 1 µM Nab2, compared to ∼42 adenosines in a reaction without rapamycin (**Fig. 4D**). The rapamycin-dependent changes in dimerization and polyadenylation were observed only when both fusion proteins were present, and rapamycin itself did not affect the polyadenylation activity of the CPAC (**Supplemental Fig. S4A-B**).

These results therefore demonstrate that the promotion of intermolecular Nab2-Nab2 interactions enhances inhibition of polyadenylation. Curiously, the concentration of Nab2 required for termination did not change noticeably whether or not Nab2-FKBP12 and Nab2-FRB were linked together before the reaction. Taking into consideration the cooperativity of Nab2 binding to poly(A) RNA (Fig. 3A), this suggests a model where two Nab2 molecules initially encounter the poly(A) RNA as monomers and subsequently interact to form a more stable dimeric assembly that terminates polyadenylation.

### Nab2 CCCH-zinc fingers 5-7 mediate poly(A) length sensing, dimerization and termination of poly(A) tail synthesis

In order to dissect the molecular basis for poly(A) tail termination activity, we next characterized the effects of Nab2 mutations employing a combination of *in vivo* and *in vitro* approaches. Poly(A) tail length control was examined *in vivo* using our previously established assay that decouples nuclear polyadenylation from cytoplasmic deadenylation by blocking mRNA export in the Mex67 anchor away (*MEX67-AA*) strain background (**Fig. 5A**, *left*). The export block sequesters endogenous Nab2 on nuclear-restricted poly(A) tails, preventing Nab2-mediated poly(A) tail length control. However, length control can be restored by over-expressing exogenous Nab2 (Turtola et al. 2021). Accordingly, detection of poly(A) tail lengths, using RNaseH digestion and Northern blotting analysis, showed that the heat-inducible *HSP104* mRNA carried hyperadenylated poly(A) tails (>70 adenosines) in nuclear export blocked cells, but these were restricted to ∼60 adenosines when exogenous Nab2 was overexpressed (**Fig. 5B**, compare ′empty′ and ′*NAB2*′). Besides controlling poly(A) tail length, Nab2 protects newly synthesized RNAs from nuclear degradation (Schmid et al. 2015; Tudek et al. 2018), as demonstrated by the higher abundance of *HSP104* mRNAs in cells over-expressing full-length Nab2.

**Figure 5.**
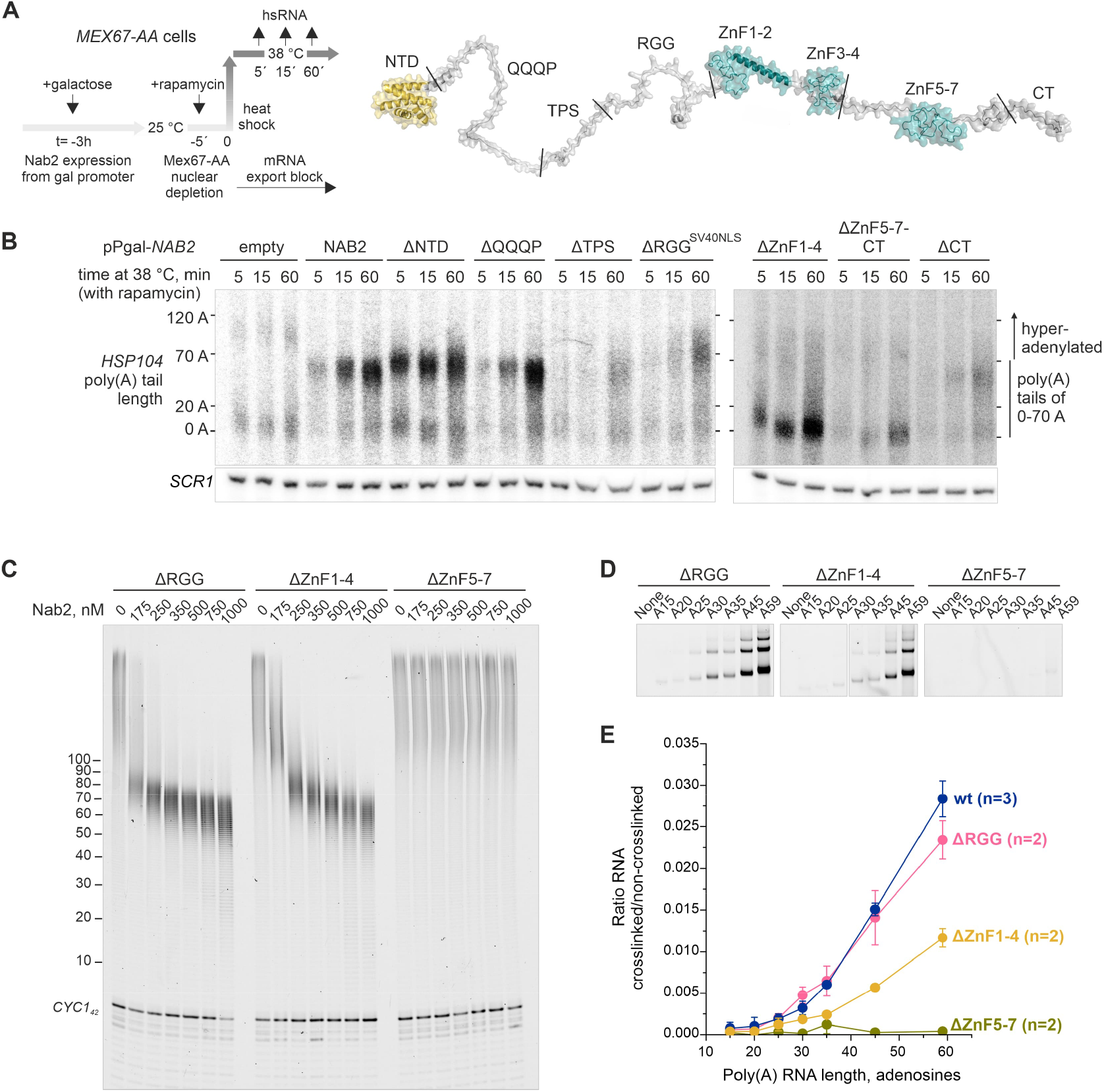
CCCH-zinc fingers 5-7 are essential for poly(A) tail length sensing. (*A*) Experimental scheme to measure nuclear mRNA poly(A) tail lengths in cells expressing mutant variants of Nab2. *Left:* mRNA export was blocked in *MEX67-AA* cells with the addition of rapamycin 5 min before transferring cells to 38 °C. RNA samples were collected at the indicated times after heat treatment. Nab2 was over-expressed for three hours prior to the heat treatment. *Right:* AlphaFold-model (Varadi et al. 2022) of Nab2 (stretched for better visualization) showing the positions of protein truncations. The regions predicted to be unstructured are shown in grey, whereas the structured NTD and ZnF are highlighted in yellow and teal, respectively. (*B*) RNaseH/Northern analysis of *HSP104* RNA 3′ends from cells subjected to different 38 °C incubation periods, depleted of nuclear Mex67, and expressing endogenous levels of Nab2 (“pPgal-empty”) or overexpressing full-length *NAB2* and mutant variants from a plasmid under control of a galactose inducible promoter (“pPgal-*NAB2*”) as indicated. See **Supplemental Fig. S5A** for the expression levels of Nab2 proteins. *HSP104* RNA was detected using an oligonucleotide probe close to the PAS, and to increase resolution, transcripts were internally cleaved with RNaseH targeted by an antisense oligo annealing 230 nucleotides upstream of the PAS. Migration of *HSP104* RNA harbouring normal (0–70 As) and hyperadenylated (>70 As) poly(A) tails are indicated. *SCR1* RNA was probed as a loading control. (*C*) CPAC-mediated *CYC1*_*42*_ polyadenylation reactions in the presence of Nab2 truncation mutants. **See Supplemental Fig. S2** for the purified proteins. (*D*) Formaldehyde-mediated RNA-protein crosslinking between Nab2 variants (500 nM) and Atto680-labelled poly(A) RNAs (100 nM) of varying lengths. Samples were run across two gels and the panels are cropped from a common exposure of those gels. The Atto680 scans of non-crosslinked RNAs and silver stained gel panels are shown in **Supplemental Fig. S5C**. (*E*) Quantification of poly(A) RNA-Nab2 crosslinks from *D*. The full-length Nab2 (wt, blue) data is replicated here from Fig. 3E.

Nab2 consists of an N-terminal-domain (NTD), analogous to a helical PWI-fold, and seven CCCH-ZnF motifs, with ZnF1-2 and ZnF3-4 folding into tandem ZnF domains, and ZnF5-7 forming an ordered domain (**Fig. 5A**, *right*). The unstructured sequence between the NTD and ZnF1 can be divided into a glutamine-proline-rich region (QQQP), the sequence surrounding the Slt2 phosphorylation site (TPS) and the arginine-glycine-glycine-motif containing domain (RGG). Finally, Nab2 harbours an unstructured C-terminal end (CT) (Truant et al. 1998; Marfatia et al. 2003; Grant et al. 2008; Martínez-Lumbreras et al. 2013; Brockmann et al. 2012; Carmody et al. 2010). To reveal which parts of Nab2 are required for poly(A) tail length control, we employed different truncated Nab2 variants. Cells expressing ΔNTD, ΔQQQP, ΔTPS, or ΔCT Nab2 variants produced *HSP104* mRNAs with poly(A) tail lengths that were similar to those from cells expressing the full-length Nab2 (**Fig. 5B, Supplemental Fig. S5A**). In contrast, poly(A) tail lengths were not restricted in cells that expressed the ΔRGG^SV40NLS^ (SV40 nuclear localisation signal (NLS) inserted at the N-terminus to compensate for the lack of NLS in the ΔRGG protein), ΔZnF1-4 or ΔZnF5-7-CT Nab2 variants.

To characterize their length control properties *in vitro*, we purified the ΔRGG, ΔZnF1-4 and ΔZnF5-7 Nab2 variants (**Supplemental Fig. S2**). Although high protein concentrations were required and poly(A) tails remained longer, the ΔRGG and ΔZnF1-4 proteins were capable of terminating polyadenylation *in vitro* (**Fig. 5C**). In contrast, the ΔZnF5-7 variant lacked the ability to terminate polyadenylation at all tested concentrations. Furthermore, this protein did not form formaldehyde-mediated RNA-protein or dimeric protein-protein crosslinks with any of the tested poly(A) RNAs (**Fig. 5D, Supplemental Fig. S5C**). While the crosslinking profile of the ΔRGG variant was similar to that of full-length Nab2, the ΔZnF1-4 variant displayed reduced efficiency. However, ΔZnF1-4 crosslinking showed unaltered poly(A) RNA length dependence with a cut-off at 25 adenosines (**Fig. 5E**).

We conclude that while the ZnF1-4 and RGG regions promote the association of Nab2 with the poly(A) tail during polyadenylation, the ZnF5-7 domain senses the minimal poly(A) tail length leading to Nab2 dimerization, in turn terminating poly(A) tail synthesis.

### Cooperation of CCCH-zinc fingers is required for Nab2 functions

A key role of ZnF5-7 in poly(A) RNA binding and poly(A) tail length control is consistent with previous studies (Kelly et al. 2010; Viphakone et al. 2008; Brockmann et al. 2012). As shown by Aibara *et al*.(Aibara et al. 2017), the isolated ZnF5-7 region homodimerizes in the presence of an A_11_G RNA, creating two binding sites for poly(A) RNA at the opposite sides of the dimer. We hypothesized that the dimeric structure might explain the observed minimal Nab2 binding length of 25 adenosines because the poly(A) RNA chain would have to be of a sufficient length to wrap around the dimer and to reach both binding sites (**Supplemental Fig. S6A)**.

To address this idea, we generated a double mutant H434D/N466D (henceforth called the dimer interface mutant) to introduce electrostatic repulsion at the dimer interface without directly affecting Nab2:poly(A) RNA interactions (**Fig. 6A**). As a control, we also created a F450A mutant to directly disrupt poly(A) RNA binding to ZnF6 (Brockmann et al. 2012). The dimer interface and the F450 mutant proteins were able to terminate polyadenylation, but resulting in slightly longer poly(A) tails than wild-type Nab2 *in vitro* (**Fig. 6B, Supplemental Fig. S6B**) and *in vivo* when observed after 5 minutes of *HSP104* induction (**Fig. 6C**). However, the most notable effect *in vivo* was the shortening of poly(A) tails of the nuclear-restricted *HSP104* mRNAs within 15-60 minutes after heat shock. These effects became even more visible when the dimer interface was further destabilised by additional substitutions (**Supplemental Fig. S6D**). Moreover, formaldehyde crosslinking indicated that both sets of mutations reduced the binding of Nab2 to poly(A) RNA (**Supplemental Fig. S6E-F**). The RNA-protein crosslinking profile of dimer interface mutant showed reduced amount of the dimeric form compared to wild-type Nab2 (**Supplemental Fig. S6G**). Thus, intermolecular protein contacts stabilize the Nab2:poly(A) RNP.

**Figure 6.**
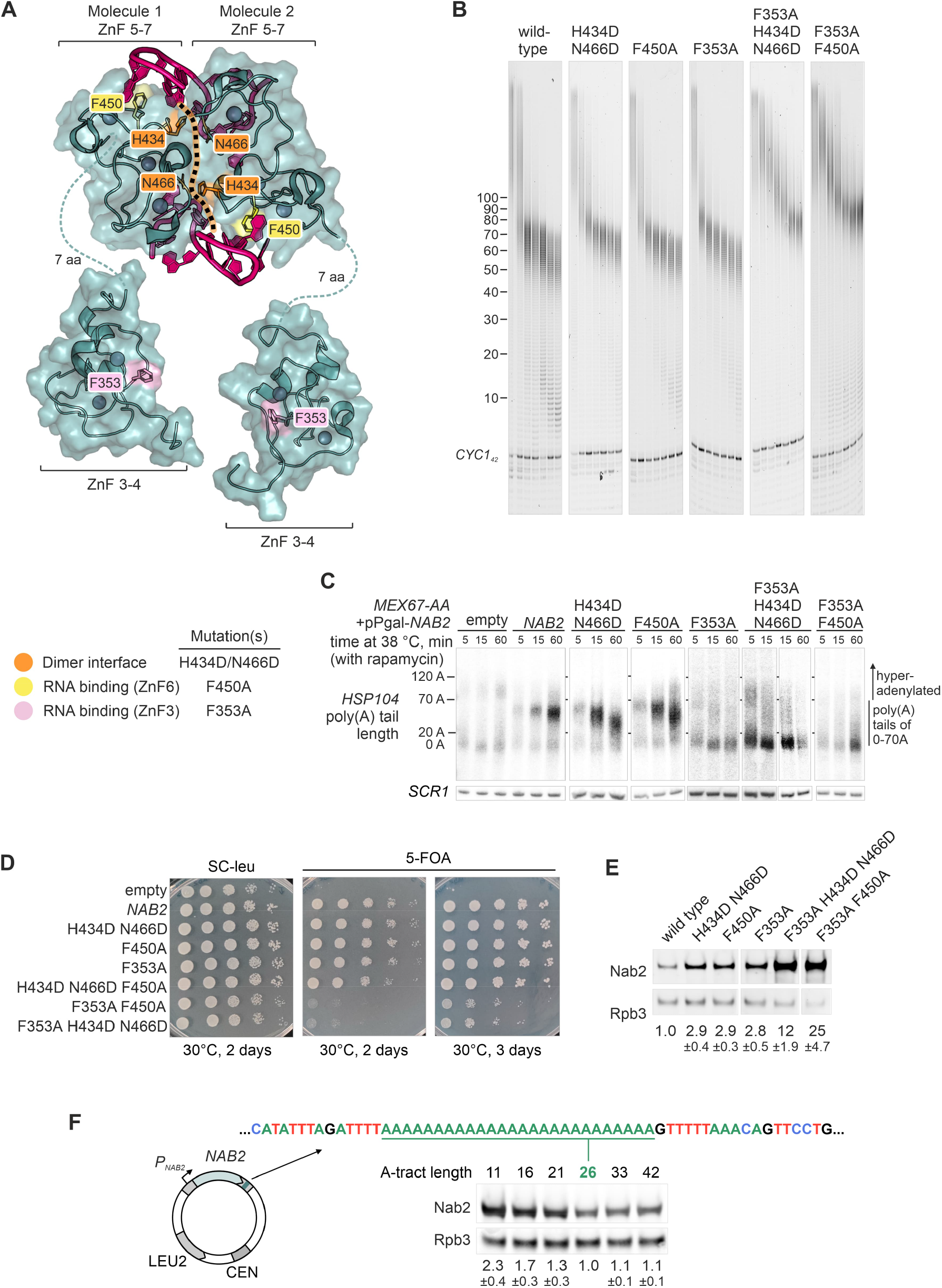
Nab2 function is tuned by the cooperation between ZnFs and the autoregulatory control of protein levels. (*A*) A composite model of the Nab2 ZnF5-7:A_11_G heterotetramer (PDB 5L2L; (Aibara et al. 2017)) and two ZnF3-4 modules (PDB 3ZJ2; (Martínez-Lumbreras et al. 2013)). The protein-protein dimer interface is indicated with a dotted line and the mutated dimer interface residues H434 and N466 are highlighted in orange; the mutated residues F450 and F353 interacting directly with RNA are highlighted in yellow and pink, respectively. (*B*) CPAC-mediated *CYC1*_*42*_ polyadenylation reactions in the presence of Nab2 wild-type and point mutant proteins. In each panel, Nab2 concentrations were: 0, 175, 250, 350, 500, 750, 1000 nM. See **Supplemental Fig. S2** for the purified proteins and **Supplemental Fig. S6B** for the side by side comparison of 1000 nM samples. (*C*) Nuclear *HSP104* mRNA poly(A) tail lengths in cells expressing mutant variants of Nab2. See Fig. 5A-B for experimental details. Nab2 expression levels are shown in **Supplemental Fig. S6C**. (*D*) Spot dilution growth assays for the *Δnab2::HIS3* p(*NAB2/CEN, URA3*) strain transformed with the p(*NAB2*/*CEN, LEU2*) plasmids expressing wild-type or mutant *NAB2* genes (or no *NAB2*; ‘empty’) under the control of endogenous *P*_*NAB2*_ promoter and downstream sequences. See **Supplemental Fig. S7A** for growth at 25 and 37 °C. *E-F*) Western blot analysis of Nab2 and Rpb3 protein levels in the *Δnab2::HIS3* p(*NAB2*/*CEN, LEU2*) strains bearing the wild-type or indicated mutant *NAB2* genes (*E*) or the A-tract length variants (*F*). The mean of Nab2/Rpb3-ratios (±standard deviation from 2-5 experiments) normalized to the wild-type is shown below. The plasmid structure of the p(*NAB2*/*CEN, LEU2*) is shown with the autoregulatory downstream sequence for the wild-type A-tract (A26^WT^; underlined). The panels in (*E*) are cropped from the same gel.

The limited effects of the dimer interface or direct RNA binding mutants on poly(A) tail length control led us to consider regions outside of ZnF5-7 as being important for termination activity. Based on results from Fig. 5, we further dissected the functions of individual ZnF1-4 and the connecting linkers between ZnF2 and ZnF3 (L2-3) and between ZnF4 and ZnF5 (L4-5). Assessment of poly(A) tails on induced *HSP104* RNAs in cells expressing individual deletions to these structural units indicated that ZnF3 and ZnF4 were required for proper termination of polyadenylation, whereas ZnF1, ZnF2, L2-3 and L4-5 were dispensable (**Supplemental Fig. S5A-B**). ZnF3 and ZnF4 fold into one module, with mainly ZnF3 predicted to contribute to direct RNA interactions (Martínez-Lumbreras et al. 2013). We thus mutated residues in ZnF3 that were previously implicated in poly(A) RNA binding (Martínez-Lumbreras et al. 2013). The F353A mutant (see Fig. 6A) conferred defects in poly(A) tail length control both *in vitro* and *in vivo* (**Fig. 6B-C, Supplemental Fig. S6B-D**), demonstrating the need for ZnF3 for timely termination of polyadenylation.

Notably, when the ZnF3 mutation F353A was combined with the dimer interface mutant or the ZnF6 mutation F450, substantially longer poly(A) tails were produced *in vitro* and higher concentrations of proteins were required for termination activity as compared to the individual ZnF domain mutants (**Fig. 6B**). Consistently, the multi-site mutants displayed pronounced defects *in vivo* (**Fig. 6C**). Similar results were obtained by introducing alternative point mutations that disrupt the RNA binding of ZnF3 (Martínez-Lumbreras et al. 2013) (**Supplemental Fig. S6A, C-D**).

Collectively, these results demonstrate that proper function of Nab2 requires the cooperation between different ZnF modules, each providing interactions that stabilize poly(A) RNA binding, either by directly contacting RNA or indirectly by stabilizing the protein-protein interface of the dimer.

### Autoregulation of *NAB2* calibrates Nab2 protein levels in a poly(A) RNA binding-dependent manner

Having established the functional importance of the intra- and intermolecular cooperation of Nab2 for its functions, we next assessed how the impairment of these different RNA-binding modes impacts cell viability. To this end, wild-type or point-mutant *NAB2* gene variants were expressed, via single-copy plasmids, under the control of endogenous upstream and downstream sequences, including the *NAB2* promoter and 3′UTR sequences. The variant-expressing plasmids were introduced into a *Δnab2* strain, maintained by the p(*NAB2*/*CEN, URA3*) plasmid, which could be counterselected on media containing 5-FOA. In line with the *NAB2* gene being essential (Hector et al. 2002; Marfatia et al. 2003), the *Δnab2* strain containing an empty p(*CEN, LEU2*) plasmid was unable to grow on 5-FOA (**Fig. 6D, Supplemental Fig. S7A**). Separate mutations targeting the dimer interface or RNA binding were well tolerated as they supported growth to the same extent as wild-type *NAB2*. However, the strains bearing multi-site mutations in both ZnF3 and ZnF5-7 domains grew noticeably slower at 25, 30 and 37 °C.

Because the concentration of Nab2 was critical for its control of polyadenylation (Fig. 2) and because the concentration requirements of the mutant proteins differed from the wild-type (Fig. 6B), we used western blotting analysis to estimate amounts of the mutated Nab2 proteins. Strikingly, protein levels were highly elevated for all of the Nab2 variants (**Fig. 6E**). Specifically, when normalized to Rpb3 levels, the ZnF3 and ZnF5-7 mutant variants displayed a 3-4-fold increased expression. Furthermore, the multi-site mutants showed 12-25-fold higher Nab2/Rpb3-ratios (**Fig. 6E, Supplemental Fig. S7B**).

This higher expression of the mutant proteins could be reconciled with previously reported Nab2 autoregulation, which is mediated by a stretch of genomically encoded adenosines (A_26_) in the *NAB2* pre-mRNA to which Nab2 can bind and interrupt the production of mature mRNA (Roth et al. 2005, 2009; Ghazal et al. 2009) (see Discussion). We shortened the A-tract from 26 to 21, 16 or 11 adenosines and observed increased expression of wild-type Nab2 protein by 1.3, 1.7 or 2.3-fold, respectively (**Fig. 6F**). Lengthening the A-tract to 33 or 42 adenosines did not significantly affect Nab2 levels. Therefore, reducing the affinity of wild-type Nab2 to the autoregulatory A-tract mimics the effects of mutant Nab2 proteins with impaired poly(A) RNA binding. Increased cellular concentrations of these supposedly compensate for their reduced ability to interact with poly(A), thereby restoring function completely, as in the case of the separate ZnF mutants, or partially, as in the case of the multi-site mutants. We conclude that the autoregulation of *NAB2* balances the poly(A) binding capacity of Nab2, thereby providing a robust control of poly(A) tail synthesis and nuclear mRNA stability.

## DISCUSSION

### A kinetic ruler controls nuclear poly(A) tail length

Tracking RNA polyadenylation reactions with high time resolution and by linking the biochemical properties of Nab2:poly(A) RNPs to reaction termination kinetics, we uncover the molecular basis of mRNA poly(A) tail length control in *S. cerevisiae*. Termination of polyadenylation by Nab2 requires its dimerization on the nascent poly(A) RNA, which can occur once the poly(A) tail is at least 25 adenosines long. This explains how Nab2 avoids terminating polyadenylation prematurely. The formation of the poly(A):Nab2 RNP likely requires that the RNA chain wraps around the dimer formed by the CCCH Zn domains 5-7, which might induce a conformational change or steric block to inhibit further polyadenylation. The minimal length for Nab2 action is consistent with a non-discrete nuclease-protected footprint of poly(A) RNA-bound Nab2 that peaks at ∼23-25 nucleotides (Viphakone et al. 2008) and with the crystal structure of dimeric Nab2 ZnF5-7 bound to A_11_G RNA (Aibara et al. 2017). The high cooperativity of binding is explained by ZnF-RNA interactions that are maximised within the dimeric ZnF5-7 unit and stabilized by the protein-protein contacts at the dimer interface. In addition to the dimerization unit, multivalent interactions through the tandem CCCH ZnF modules and the RGG unit facilitate timely termination. These regions may promote the productive Nab2:poly(A) RNP assembly pathway by increasing the association rates and the RNA-residence times of monomeric proteins prior to the formation of cooperative dimeric interactions.

Importantly, neither the minimal RNA-footprint nor the binding architecture of Nab2 are sufficient to explain how the protein restricts poly(A) tail lengths to ∼60 adenosines. We solve this conundrum by considering polyadenylation kinetics, demonstrating that poly(A) tail length control is a non-equilibrium process. Eventual tail length is an outcome of the relative rates of polyadenylation by the CPAC and Nab2-poly(A) RNA association. As a result, poly(A) tail length is not “measured” *per se*, rather the on-rate of Nab2 binding to the poly(A) tail decides its length. The on-rate increases with increasing tail length, and is proportional to the concentration of Nab2. We therefore propose that Nab2 is a “kinetic ruler”, the concentration of which is used for quantifying poly(A) tail length (**Fig. 7**).

**Figure 7.**
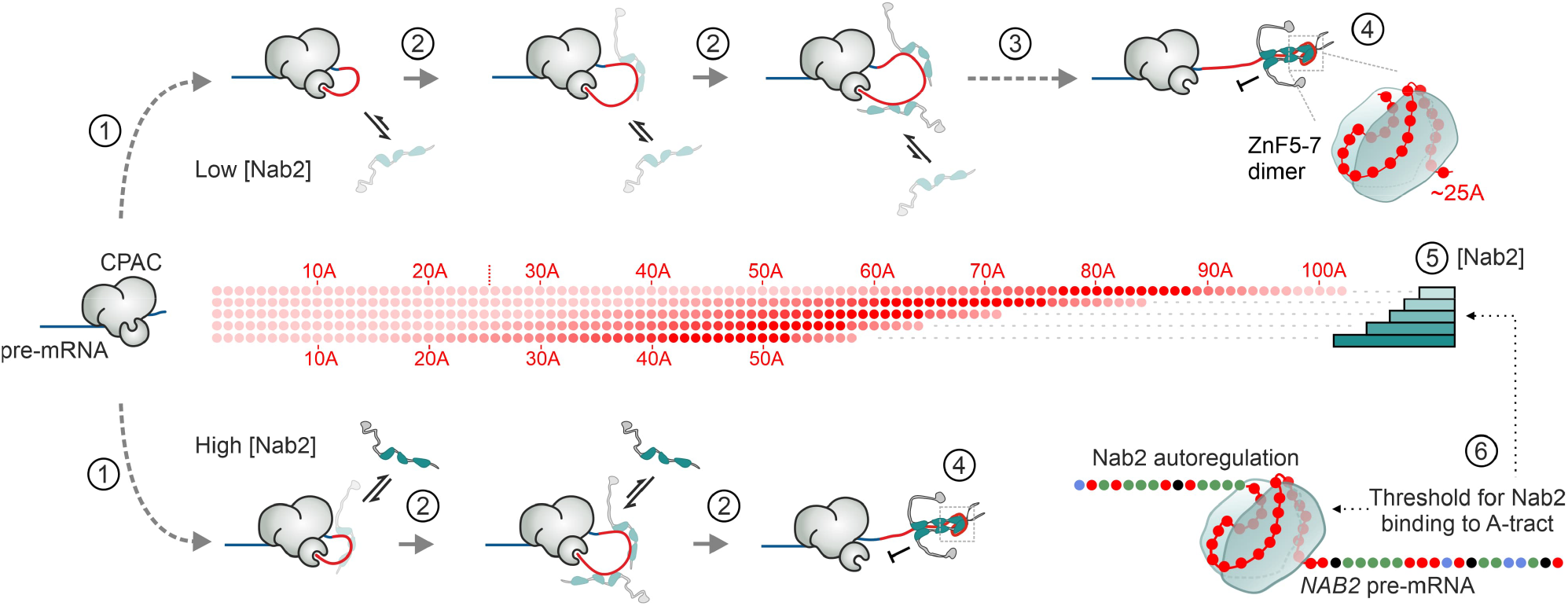
Kinetic ruler mechanism of poly(A) tail length control. The control of pre-mRNA (*blue line*) polyadenylation is depicted with Nab2 (*grey/teal*) present at low or high concentration (*top and bottom rows*, respectively). *1*) The poly(A) tail (*red lines*) is synthesized by the CPAC (*grey*) that binds to the PAS and tethers the poly(A) polymerase to the pre-mRNA. The start of poly(A) tail elongation is limited by a slow step of unknown nature (*curved dashed arrows*). *2*) The first 70 adenosines of the poly(A) tail are elongated within three seconds (*short grey arrows*). The rate of Nab2 binding to the RNA (*black small arrows*) increases whilst the poly(A) tail grows longer. *3*) The polyadenylation activity of the CPAC decreases after the synthesis of ∼70 adenosines (*long dashed arrow*). *4*) Dimerization of Nab2 on poly(A) tail, mediated by ZnF5-7 and stabilized by other ZnF domains, terminates polyadenylation. Dimerization is limited to poly(A) RNAs that are longer than 25 adenosines. *5*) Concentration of Nab2 determines the length of poly(A) tails (*individual adenosines depicted as red spheres*): at low concentrations (*bars, light teal*) poly(A) tails become long whereas at high concentrations (*bars, dark teal*) shorter poly(A) tails are produced. *6*) Concentration of Nab2 is autoregulated through a A_26_-sequence within the *NAB2* pre-mRNA. The binding properties of Nab2 to this sequence locks its nuclear-available concentration.

### Calibration and tuneability of the kinetic ruler

While the suggested kinetic ruler provides a solution to the problem of RNA chain length-sensing, such a fine-tuned mechanism requires careful titration of Nab2 protein concentrations in order to maintain the production of mRNAs with specific poly(A) tail lengths. This is ingeniously managed by an autoregulatory system where Nab2 negatively impacts the production of its own mRNA. Here, binding of Nab2 to an encoded A_26_-tract within the *NAB2* pre-mRNA prevents the co-transcriptional utilisation of the PAS by the CPAC. This leads to an alternative RNase III-triggered transcription termination event, which in turn elicits exosome-mediated degradation of the *NAB2* pre-mRNA. Conversely, if Nab2 is not available, the CPAC cleaves the pre-mRNA upstream of the A_26_-sequence and produces a mature *NAB2* mRNA (Roth et al. 2005, 2009; Ghazal et al. 2009).

According to the principles of autoregulatory gene networks (Alon 2007), the binding affinity of Nab2 to the A_26_-tract determines and locks the steady-state level of nuclear-available Nab2. Consistently, we observed drastic upregulation of the levels of mutant Nab2 proteins with reduced ability to bind poly(A) RNA. Based on our results, the A_26_-tract is sufficient to allow Nab2 dimerization and has apparently evolved to specifically calibrate the poly(A) binding capacity of Nab2 by probing dimer formation (**Fig. 7**). Interestingly, long A-tracts are found downstream of *NAB2* genes in various fungal genomes, often mixed with non-A nucleotides. For example, *Candida auris*, an emerging fungal pathogen, has 51 adenosines in a 57-nucleotide region. The implication from the kinetic ruler mechanism, and autoregulated PABP levels, is that global mRNA poly(A) tail lengths may be controlled by these autoregulatory A-tracts.

Nab2 protein levels might still fluctuate on timescales shorter than the response time to establish a new equilibrium, or display a non-uniform distribution within the nucleus, with consequences for mRNA production. For example, Nab2 becomes sequestered into foci following nuclear accumulation of poly(A) RNA in conditions such as heat shock (Carmody et al. 2010) and glucose depletion (Heinrich et al. 2024), or as a result of defects in several RNA processing pathways that impair mRNA export (Paul and Montpetit 2016; Aguilar et al. 2020; Tudek et al. 2018). The kinetic ruler model now provides a rationale for the hyperadenylation phenotype observed in yeast strains impaired in mRNP processing and export (Hilleren and Parker 2001; Jensen et al. 2001; Libri et al. 2002; Rougemaille et al. 2007), and predicts how changes in Nab2 availability impact mRNA poly(A) tail lengths and nuclear stabilities. Different domains of Nab2, including the unstructured QQQP, TPS and RGG regions influence its biomolecular clustering properties, nuclear-cytoplasmic shuttling behaviour and interactions with mRNP proteins (Heinrich et al. 2024; Marfatia et al. 2003; Iglesias et al. 2010), and these may be subjected to regulation via post-translational modifications (Green et al. 2002; Carmody et al. 2010; Insco et al. 2023). An intriguing question for future studies is therefore whether Nab2 modifications may re-calibrate the kinetic ruler to alter poly(A) tail lengths, thereby tuning regulatory properties of individual RNAs or across the entire transcriptome.

### Distinct poly(A) tail elongation and termination mechanisms in yeast and mammals

Despite the high level of conservation of the CPAC-machineries across eukaryotes, poly(A) tails are elongated in distinctively different manners in yeast and mammalian cells. A major difference is the basal polyadenylation processivities: yeast Pap1 is tightly bound to CPAC and synthesizes poly(A) tails efficiently up to ∼70 adenosines (this study), whereas the mammalian PAP, which is more loosely associated with the CPAC machinery, requires additional stimulation by PABPN1 during poly(A) tail elongation (Rodríguez-Molina and Turtola 2023; Eckmann et al. 2011; Kerwitz et al. 2003). Secondly, the kinetic ruler model of tail length control in yeast fundamentally differs from the proposed model in mammals. In the current view, mammalian mRNA poly(A) tail length is determined by a molecular ruler mechanism whereby ∼15-20 molecules of PABPN1 cover the ∼250 adenosine long poly(A) tail and terminates the processive phase of tail elongation by obstructing interactions between PAP, PABPN1 and the rest of the CPAC machinery (Kühn et al. 2009; Keller et al. 2000). However, this picture of mammalian polyadenylation control is confounded by observations from several organisms, indicating that the depletion of Nab2/ZC3H14 leads to increased lengths of bulk poly(A) tails (Pak et al. 2011; Kelly et al. 2014; Rha et al. 2017; Bienkowski et al. 2017; Morris and Corbett 2018; Li et al. 2024). This raises the possibility that Nab2/ZC3H14 may regulate the timing of poly(A) tail termination alongside PABPN1. Notably, the human ZC3H14 and yeast Nab2 appear to share the RNA binding modality needed for termination as over-expression of a chimeric protein, where ZnF1-7 of yeast Nab2 were replaced by ZnF1-5 of human ZC3H14, controlled poly(A) tail length and protected the nuclear-restricted *HSP104* mRNA in yeast (**Supplemental Fig. S5A & S5D**). Further studies are needed to clarify the roles and possible interplay of different factors in this process.

### Architectural and kinetic considerations of PABP-mediated regulation of RNA processing

Poly(A) tail-bound nuclear PABPs form protective structures at RNA 3′ends and promote mRNA export, yet they can also act within different adaptor complexes to target various transcript classes for decay, but the logic behind these choices remains largely unknown (reviewed in (Rambout and Maquat 2024)). Nab2/ZC3H14 has been mostly linked to protective functions (Grenier St-Sauveur et al. 2013; Latour et al. 2025; Morris and Corbett 2018; Tudek et al. 2018; Schmid et al. 2015; Turtola et al. 2021), whereas PABPN1 works in several decay pathways (Meola et al. 2016; Bresson et al. 2015; Latour et al. 2025). The characterization of Nab2 in this study suggests profound differences in the biochemical properties of ZC3H14 compared to those of PABPN1 that might provide clues to their antagonistic functions.

First, our mutagenesis revealed the requirement of Nab2 dimerization and multidomain interactions in protecting the 3′ends of long poly(A) tails against polymerase and nuclease activities, thus, showing how it preserves poly(A) tail lengths in the nucleus. In contrast, PABPN1 leaves the RNA 3′ends accessible for polymerase and nuclease activities (Nguyen et al. 2015; Kerwitz et al. 2003). Second, Nab2 proteins bind poly(A) RNA in a cooperative manner, conferring a preference to bind >25 adenosine tails. PABPN1, instead, binds 11 adenosines without notable cooperativity emerging for longer poly(A) RNA substrates (Meyer et al. 2002). Therefore, poly(A) tail length-dependent recruitment of different PABPs might be one mechanism to distinguish between transcript types and to direct them to different fates.

Echoing the tuneable kinetic ruler mechanism proposed here, the coupling of poly(A) tail length to translational efficiency in oocytes and early embryos requires a concentration regime of PABPC that favours its association with long-tailed mRNAs (Xiang and Bartel 2021). Intriguingly, Pab1 (the yeast homologue of PABPC) can also inhibit polyadenylation and control poly(A) tail lengths in a concentration dependent manner (Turtola et al. 2021; Viphakone et al. 2008), raising the possibility that the length control principles uncovered here could play a role in the complex dynamics of cytoplasmic polyadenylation and deadenylation (Xiang et al. 2024; Lim et al. 2016). Our results therefore highlight the need to understand both structural and kinetic properties behind PABP-mediated regulatory mechanisms.

## MATERIALS AND METHODS

### DNA constructs

The cloning involving the pESC-URA (pPgal), p(*CEN, LEU2*) and pET28b was performed in DH5α or XL1 *E. coli* cells. A list of plasmids is in **Supplemental Table S2**. *NAB2* deletions and point mutations were first constructed in pESC-URA vectors. The N-terminal (ΔNTD) and C-terminal (ΔZnF5-7-CT and ΔCT) truncations were created by amplifying the included fragment of the coding region by PCR with primers containing overhangs with EcoRI sequence at the 5′end and the NotI site at the 3′end of the amplified sequence (see **Supplemental Table S3** for primer sequences and DNA templates used in the PCR reactions). The internal deletions were constructed in two rounds of PCR by first amplifying the N-terminal and C-terminal fragments separately with primers containing overhangs that created complementary sequences across the deleted region. These purified fragments were then used as a template in a second PCR reaction with primers that annealed to the N-terminal and C-terminal ends and contained overhangs with EcoRI and NotI sites, respectively. The ΔQQQP fragment was amplified using pAC1039 (Marfatia et al. 2003) as a template in PCR. The chimeric gene where ZnF1-7-CT region of *S. cerevisiae* Nab2 was replaced with ZnF1-5 of human ZC3H14 was constructed by fusing two fragments in a PCR reaction: a PCR-amplified N-terminal fragment of Nab2 (amino acid residues 1-255), and a DNA string fragment (Thermo Scientific/Gene Art) of human ZC3H14 (amino acid residues 594-736; codon optimized for expression in *S. cerevisiae*). The PCR products were cloned into the pESC-URA vector under the pGal10 promoter using EcoRI and NotI. In the indicated constructs (**Supplemental Table S2**) the fragment was cloned in-frame with the C-terminal Flag-tag sequence by omitting the stop codon from the 3′end overhang of the amplified sequence. *NAB2* point mutations were created by inverse PCR, amplifying the whole plasmid DNA, followed by DpnI digestion of the template plasmid, T4PNK phosphorylation of linear DNA ends, and blunt end ligation. The *NAB2* coding sequences were confirmed by Sanger DNA sequencing and the overall composition of the plasmids were analysed by restriction digestion. The p(*CEN, LEU2*) parent plasmid was constructed by PCR-amplifying sequences 400 bp upstream and 400 bp downstream of the *NAB2* ORF with complementary overhangs using pAC1039 (Marfatia et al. 2003) as a DNA template, and inserted into HindIII and SacI-linearized pRSII415 vector (Chee and Haase 2012) through NEBuilder HiFi DNA Assembly (New England Biolabs). Subsequently, p(*NAB2*/*CEN, LEU2*) plasmids were generated by cloning the *NAB2* genes (digested from the pESC-URA plasmids) using the EcoRI and NotI sites between the upstream and downstream sequences. The A-tract variants were created by inverse PCR, as indicated above, using the p(*CEN, LEU2*) parent plasmid as a template. The NdeI-SacI fragments were then cloned into the p(*NAB2*/*CEN, LEU2*) plasmid to replace the wild-type autoregulatory sequence with variable A-tract sequences. To construct the *E. coli* expression vectors, *NAB2* genes, originally cloned into the pESC-URA plasmids (see above), were PCR-amplified using primers with specific overhangs. The resulting PCR products were inserted into NcoI and NotI -linearized pET28b vectors through NEBuilder HiFi DNA Assembly. To construct the Nab2-FKBP12 and Nab2-FRB fusion proteins, the PCR-amplified fragments of human FKBP12 (amino acid residues 1-107) or FRB (amino acid residues 2021-2113 of mTOR, T2098L) containing appropriate overhangs were included in the assembly reactions. The pEG012 (ΔZnF5-7) was constructed by inverse PCR using pEG005 (Nab2 wild-type; pET28b) as a template and primers that created the deletion (amino acids 390-485 deleted), followed by DpnI digestion of the template plasmid, T4PNK phosphorylation of linear DNA ends, and blunt end ligation.

### Expression of CPF modules, CF IA and SII-Nab2 in insect cells

The *PFS2* and *REF2* genes encoded a C-terminal Twin-Strep (SII)-tag. *PCF11* gene encoded an N-terminal SII-tag and *RNA14* encoded an N-terminal 8xHis-tag. *NAB2* gene encoded an N-terminal SII-tag. Vectors (see **Supplemental Table S2**) carrying the polymerase module genes (*CFT1, YTH1, FIP1, PAP1, PFS2*-SII) (Hill et al. 2019), nuclease module genes (*YSH1, CFT2, MPE1*) (Kumar et al. 2021), phosphatase module genes (*PTA1, PTI1, SSU72, SWD2, GLC7, REF2*-SII) (Kumar et al. 2021), CF IA genes (8xHis-Rna14, Rna15, SII-Pcf11, Clp1) (Kumar et al. 2021), or SII-*NAB2* (Turtola et al. 2021) gene were transformed into chemically competent DH10EmbacY cells and bacmids were isolated as previously described (Bieniossek et al. 2008). *Spodoptera frugiperda Sf9* cell line was used for the expression of baculovirus-based recombinant proteins and complexes. For CPF modules and CF IA, the bacmid transfection (production of P1 virus) was performed in *Sf9* cells that were cultured to log phase in Sf-900 II SFM medium (Gibco) before being transferred to ExpiSF CD Medium (Gibco) at a density of 0.5 × 10^6^ cells/ml. Fugene HD transfection reagent was used following the manufacturer’s protocol (Promega). The cultures were incubated at 27 °C and monitored daily for signs of infection. The supernatant containing the P1 virus was collected, sterile-filtered, and stored at 4 °C with the addition of 2% FBS for stabilization. To amplify the P2 virus, a 1:75 (v/v) dilution of the P1 virus was used to infect suspension culture of *Sf9* cells, which had been grown in Sf-900 II SFM and then transferred to ExpiSF medium at a density of 1.5 × 10^6^ cells/ml. The infected cells were cultured at 27 °C 140 rpm for 72-96 hours before harvesting the P2 virus stock. Large-scale infections for protein expression were carried out in 500 ml cultures of *Sf9* cells at 2 × 10^6^ cells/ml in Sf-900 II SFM or ExpiSF medium (27 °C, 140 rpm, 72-96 h) with 1:50 to 1:100 (v/v) P2 virus. The P2 viruses for nuclease module (1:60) and phosphatase module (1:75) were used for co-infection to express both modules simultaneously. Cells were harvested by centrifugation at 2,500 × g for 15 min and washed in PBS. Pellets were flash frozen in liquid nitrogen and stored at -80 °C. SII-Nab2 was expressed as described in (Turtola et al. 2021).

### Expression of CF IB (6xHis-Hrp1) and Nab2-6xHis in E. coli

CF IB (6xHis-Hrp1) (Casañal et al. 2017) and Nab2-6xHis proteins were expressed in Xjb RIL *E. coli* cells transformed with plasmids detailed in **Supplemental Table S2**. The cells were grown in LB medium at 37°C. For Nab2-6xHis expression, the medium was supplemented with 0.1 mM ZnCl_2_. IPTG (1 mM) and arabinose (0.1%, w/v) were added at an optical density (OD_600_) of approximately 0.6, and the cells were harvested 3-5 hours later.

### Purification and reconstitution of CPF

CPF polymerase module was purified as in (Rodríguez-Molina et al. 2022). Frozen pellet from 1 l culture was thawed in lysis buffer (50 mM Hepes-NaOH, pH 8.0, 300 mM NaCl, 1 mM TCEP) supplemented with cOmplete EDTA-free protease inhibitor cocktails (Sigma-Aldrich), and 1 ml BioLock biotin blocking solution (IBA Lifesciences). Cells were lysed by sonication and lysates were cleared by centrifugation at 40 000 × g for 15 min at 4 °C. The clarified lysate was bound in batch to 3 ml Strep-Tactin Sepharose beads (IBA Lifesciences) for 1 h on a rotating platform at 4 °C and subsequently transferred to gravity-flow column. Beads were washed extensively with lysis buffer, and subsequently incubated for 20 minutes with the elution buffer (50 mM Hepes-NaOH, pH 8.0, 150 mM NaCl, 1 mM TCEP and 1.2 mg/ml desthiobiotin) before eluting. The eluted sample was filtered and loaded onto a 1 ml Hitrap Q anion exchange chromatography column that was equilibrated using buffer that contained 20 mM Hepes-NaOH, pH 8.0, 150 mM NaCl, 0.5 mM TCEP. The polymerase module was eluted using a NaCl gradient from 150 to 600 mM over 40 CV. The fractions containing all polymerase module subunits were pooled, concentrated using a 100 kDa molecular weight cut-off (MWCO) concentrators and snap-frozen in liquid N_2_. The purification of the CPF nuclease-phosphatase module was carried out as described above, with different buffers. Here, the lysis buffer contained 50 mM Hepes-KOH, pH 8.0, 150 mM KCl, 0.5 mM magnesium acetate and 0.5 mM TCEP. The anion exchange chromatography was carried out using a 5 ml ResourceQ column (Cytiva) and a lysis buffer-based NaCl gradient from 150 to 500 mM over 15 CV. The complete CPF complex was reconstituted from separately purified polymerase module and the nuclease-phosphatase modules as in (Rodríguez-Molina et al. 2022). The CPF was reconstituted at final 10 µM by mixing in 1:1 molar ratio preparations of polymerase module and nuclease-phosphatase modules and separating the mixture in a Superose 6 Increase column. Peak fractions were analysed by SDS-PAGE and the fractions containing stoichiometric ratios of individual subunits were pooled, concentrated and snap-frozen in liquid N_2_ for storage.

### Purification of CF IA

CF IA was purified according to (Kumar et al. 2021). Frozen pellet from 2 l culture (∼40 ml) was thawed into a total volume of 150 ml in lysis buffer (50 mM Hepes-NaOH, pH 7.9, 250 mM NaCl, 5% (w/v) glycerol, 0.5 mM TCEP) supplemented with 2 µg/ml DNase I (Thermo Scientific), 4 × cOmplete EDTA-free protease inhibitor cocktail tablets (Sigma-Aldrich), and 1 ml BioLock biotin blocking solution (IBA Lifesciences). Cells were lysed by sonication and lysates were cleared by ultracentrifugation at 42 000 × rpm in a Beckmann 50.2 Ti rotor for 45 min at 4 °C. The clarified lysate was bound in batch to 3 ml Strep-Tactin Sepharose beads (IBA Lifesciences) for 1 h on a rotating platform at 4 °C and subsequently transferred to a gravity-flow column. Beads were washed with 150 ml of lysis buffer, and subsequently incubated for 20 minutes with the lysis buffer supplemented with 1.2 mg/ml desthiobiotin before eluting. The protein sample was loaded onto a 5 ml HiTrap Heparin column (Cytiva), eluted using a NaCl gradient from 250 mM to 1000 mM over 15 CV, and the peak fractions were concentrated using a 10 kDa MWCO concentrator. The concentrated protein was further purified through Superdex 200 Increase size exclusion chromatography column using a buffer that contained 20 mM Hepes-NaOH, pH 8.0, 250 mM NaCl, 0.5 mM TCEP. The fractions containing all CF IA subunits were pooled and concentrated before being snap-frozen in liquid N_2_ for storage.

### Purification of CF IB (6xHis-Hrp1)

CF IB was purified as in (Casañal et al. 2017). Cell pellets were resuspended in Hrp1 lysis buffer (50 mM Hepes-KOH, pH 8.0, 300 mM NaCl, 0.5 mM TCEP, 20 mM imidazole) supplemented with 10% (w/v) glycerol, 2 µg/ml DNase I (Thermo Scientific), 2 µg/ml RNase A (Thermo Scientific) and cOmplete EDTA-free protease inhibitor cocktail tablets (Sigma-Aldrich). The cells were lysed by sonication on an ice bath, using 30-second on/off cycles for a total of 15 min. The lysate was then centrifuged at 20000 rpm for 30 min at 4 °C using a Beckmann JA-25.50 rotor. The cleared lysate was incubated with Ni-NTA beads on a rotating platform at 4 °C for 1.5 hours and subsequently transferred to gravity-flow column. The column was washed sequentially with the Hrp1 lysis buffer and wash buffer (50 mM Hepes-KOH, pH 8.0, 300 mM NaCl, 0.5 mM TCEP, 30 mM imidazole) before the Hrp1 protein was eluted with elution buffer (50 mM Hepes-KOH, pH 8.0, 300 mM NaCl, 0.5 mM TCEP, 300 mM imidazole). The eluate was directly loaded onto a 5 ml HiTrap Heparin column (Cytiva), equilibrated with a buffer containing 20 mM Hepes-KOH, pH 8.0, 100 mM NaCl, 0.5 mM TCEP. The protein was eluted using a NaCl gradient from 100 to 1000 mM over 10 CV. Fractions containing Hrp1 were pooled, concentrated with a 10 kDa MWCO concentrator (Pierce), and further purified by size exclusion chromatography in Superdex 200 Increase column (Cytiva) using a buffer that contained 20 mM Hepes-KOH, pH 8.0, 250 mM NaCl, 0.5 mM TCEP, and snap-frozen in liquid N_2_.

### Purification of Nab2 (wild-type and mutants)

SII-tagged Nab2 expressed in insect cells was purified as in (Turtola et al. 2021), with modifications. Cell pellet from 2 l culture was resuspended in lysis buffer (50 mM Tris-HCl, pH 8.0, 50 mM NaCl, 2 mM MgCl_2_, 0.1 mM ZnCl_2_, 1 mM TCEP) supplemented with 4 µg/ml DNaseI (Sigma-Aldrich), 4 µg/ml RNaseA and 42 U/ml Benzonase (Sigma-Aldrich). The cells were lysed using sonication, followed by centrifugation at 18000 rpm for 15 min at 4 °C using a Beckmann JA-25.50 rotor. The cleared lysate was supplemented with BioLock biotin blocking solution (IBA Lifesciences) and incubated for 20 additional min at 20 °C for completing nuclease digestion. NaCl concentration of the lysate was adjusted to 200 mM before incubating the lysate with the Strep-Tactin beads for 2.5 h under rotation at 4 °C. The beads were then moved to a gravity flow column and washed with wash buffer (20 mM Tris-HCl, pH 8.0, 0.1 mM ZnCl_2_, 0.5 mM TCEP) containing 200 mM NaCl and 0.2% Tween-20, then with wash buffer containing 300 mM NaCl, wash buffer with 500 mM NaCl, wash buffer with 75 mM NaCl, and finally eluted with wash buffer containing 75 mM NaCl and 1.2 mg/ml desthiobiotin. The eluate was combined with three volumes of buffer A (20 mM MES, pH 6.5, 75 mM NaCl, 0.1 mM ZnCl_2_, 0.5 mM TCEP), loaded to a ResourceS column (Cytiva) and eluted using a NaCl gradient. Peak fractions were combined and concentrated using 10 kDa MWCO concentrator before snap-freezing aliquots in liquid N_2_. To purify the Nab2-6xHis proteins expressed in *E. coli*, the cell pellets were resuspended in Nab2 lysis buffer (50 mM Hepes-KOH, pH 7.4, 500 mM NaCl, 5% (w/v) glycerol, 1 mM TCEP, 0.2% Tween-20 (v/v), 0.1 mM EDTA, 0.1 mM ZnCl_2_) supplemented with cOmplete EDTA-free protease inhibitor cocktail tablets (Sigma-Aldrich). The cells were lysed using sonication on an ice bath, applying 30-second on/off cycles for a total of 15 min. The lysate was then centrifuged at 20000 rpm for 30 min at 4 °C using a Beckmann JA-25.50 rotor. The cleared lysate was supplemented with 20 mM imidazole and loaded on a GoBio Zn-IDA column (Bio-Works) to capture the 6xHis-tagged Nab2. The column was washed with 10 column volumes (CV) of the Nab2 lysis buffer containing 20 mM imidazole, followed by 6 CV of lysis buffer containing 50 mM imidazole. The protein was then eluted using lysis buffer supplemented with 200 mM imidazole. The first 15 ml of the elution were mixed with 35 ml of Buffer A (20 mM Hepes-KOH, pH 7.4, 5% glycerol, 0.5 mM TCEP, 0.1 mM EDTA, 0.1 mM ZnCl_2_), and the mixture was filtered before loading onto to a 5 mL GoBio MiniS column (Bio-Works) for cation exchange chromatography. The column was washed with 5 CV of Buffer A, followed by a steep NaCl gradient from 150 to 1000 mM over 4.2 CV. 6xHis-Nab2 began eluting at ∼400 mM NaCl. Eight earliest fractions containing Nab2 (total 8 ml) were pooled and concentrated to 1 ml using 10 kDa MWCO concentrators (Pierce). The concentrated sample was centrifuged at 10000 × g for 10 min at 4 °C before 500 µl was injected into a Superdex 200 column (Cytiva) equilibrated with gel filtration buffer (20 mM Hepes-KOH, pH 7.4, 500 mM NaCl, 0.5 mM TCEP, 0.1 mM EDTA, 0.1 mM ZnCl_2_). Peak fractions of the gel filtration run containing Nab2, as detected by SDS-PAGE analysis, and displaying lowest A_260_/A_280_-ratios were pooled and concentrated further in 10 kDa MWCO concentrator before snap-freezing aliquots in liquid N_2_. The purification of Nab2 ΔRGG was performed otherwise as described above, with the exception that the buffers used for MiniS cation exchange and the elution step from the Zn-IDA column were prepared with MES buffer (pH 6.0) instead of Hepes buffer (pH 7.4). The purification of Nab2 ΔZnF1-4 was carried out as described above up to the MiniS cation exchange step. Due to a high nucleic acid content in the MiniS eluate, the peak fractions were pooled and diluted 1:10 in Buffer A containing 120 mM NaCl and 2 mM MgCl_2_. The diluted sample was treated overnight with 50 U/ml benzonase (Pierce) at 4°C. Afterwards, benzonase and degraded nucleic acids were removed by a second MiniS cation exchange chromatography step, performed as described above. The peak fractions were then pooled, concentrated, and snap-frozen for storage. The purification of Nab2 ΔZnF5-7, which has a lower isoelectric point than the wild-type protein, followed the same procedure as the wild-type, except that a MiniQ column (Bio-Works) was used instead of a MiniS column during the second purification step to perform anion exchange chromatography using the same buffers.

### RNA oligonucleotides

Oligonucleotide sequences and information are provided in **Supplemental Table S4**. RNA oligonucleotides were purchased from Eurogentec. Certain RNAs were prepared in-house using splinted ligation. This method employed a complementary DNA splint to align the 3′-OH and 5′-phosphate termini of two RNA strands, facilitating their ligation by T4 DNA ligase (Thermo Scientific). The reaction products were resolved on denaturing urea-polyacrylamide gels, detected through the fluorescence of 5′-Atto680 labels, excised from gel and purified using ZR small-RNA PAGE Recovery Kit (Zymo Research).

### Polyadenylation reactions

All reactions (modified from those performed in (Turtola et al. 2021)) were performed at 30 °C in polyadenylation buffer that contained 25 mM Hepes-KOH, pH 8.0, 150 mM potassium acetate, 2 mM magnesium acetate, 0.05 mM EDTA, 0.1 U/µl RiboLock RNase Inhibitor (Thermo Scientific). Most reactions were set up by pre-incubating the RNA substrate with CF IA and CF IB for 5 minutes at 30 °C in 1/3 of the final reaction volume. Another 1/3 reaction volume of solution containing CPF, and, where specified Nab2-6xHis, was added and pre-incubated for additional 3 minutes. Polyadenylation was initiated by the addition of 1/3 volume of a solution that contained ATP. Final concentrations of reaction components, the order of addition of CPAC components and the pre-incubation periods in each polyadenylation experiment are specified in **Supplemental Table S5**. Reactions were stopped after 4 minutes, or in the case of time-course experiments, at the indicated time points by the addition of 0.7 reaction volumes of 2 M HCl, and immediately neutralized by the addition of 5.3 volumes of neutralization solution (290 mM Tris base, 13 mM EDTA, 0.025% (w/v) bromophenol blue, 94% formamide). Time-resolved polyadenylation reactions were performed using the RQF 3 quench-flow instrument (KinTek Corporation, Austin, TX, USA). The reaction was initiated by rapid mixing of 16 µl of preformed CPAC with 16 µl of 4 mM ATP solution (both solutions in polyadenylation buffer). The reaction was allowed to proceed for 0.2–15 s at 30 °C and quenched with 83 µl of 0.5 M HCl, and immediately neutralized by adding 165 µl of neutralization solution. Product RNA were separated on 13% denaturing polyacrylamide gels and RNA species were visualized with Odyssey Infrared Imager (Li-Cor Biosciences) or Sapphire Biomolecular Imager (Azure Biosystems); gel lane profiles were quantified using ImageJ software. The poly(A) tail length at each time point was quantified by plotting the gel lane intensity in Image J, and analysed in Origin by subtracting the background level and integrating the signal across the region where the poly(A) tail signal appears (wave-front). The nucleotide position corresponding to the midpoint of the cumulative signal was identified, representing the mean poly(A) tail length of the elongating RNA population. Linear fitting was performed with instrumental weighing method.

### SwitchSENSE

The interactions of SII-Nab2 with RNA and DNA were analyzed on a DRX2 instrument (Dynamic Biosensors GmbH) using a MPC2-48-2-G1R1-S chip equilibrated with SwitchSENSE buffer 12.5 mM Hepes-KOH, pH 8.0, 75 mM potassium acetate, 1 mM magnesium acetate, 0.025 mM EDTA) at 25 °C. Prior to each kinetic analysis oligonucleotides cNLA-X2 with CX2 and cNLB-X1 with either CX1-rA30 or CX1-rA60 (see Supplemental Table 4), together were annealed to DNA strands attached on the chip surface (NL-A48 and NL-B48, respectively) by flowing 200 nM oligonucleotide over the chip for 4 min in a buffer of 10 mM Tris-HCl, pH 7.4, 40 mM NaCl, 0.05 % (v/v) Tween-20, 50 μM EDTA, 50 μM EGTA. As a result, 68-bp dsDNA nanolevers were formed without (DNA) or with a 30-nucleotide (rA30) or 60-nucleotide (rA60) RNA overhang extending in the 5′ to 3′ direction. SII-Nab2 in a 1:3 dilution series, starting at 100 nM was injected at 50 µl/min for between 9 min followed by dissociation in running buffer after the highest concentration for 80 minutes at 50 µl/min. Dynamic switching data were analysed using the supplied SwitchBUILD software using a 1:1 kinetic model to give value for the association rate constant, *k*_*on*_, dissociation rate constant, *k*_*off*_ and to calculate the kinetic dissociation constant *K*_*d*_ *= k*_*off*_ */k*_*on*_. Averages with standard errors were calculated for repeat experiments (n = 4 for rA30 and rA60; n = 6 for control DNA)

### Electromobility shift assay (EMSA)

A_59_ RNA (10 nM) and Nab2-6xHis (0-500 nM) were mixed in polyadenylation buffer that additionally contained 5% (v/v) glycerol and 0.025% OrangeG, and incubated at 30 °C for 1.5 h. The mix was then loaded on a pre-cooled 6% native polyacrylamide gel prepared in His-MES buffer system (38 mM L-histidine, 5.86 g/l MES hydrate, pH 6.4 determined at 25 °C). Electrophoresis was carried out at 300 V for 1h at 4 °C. The Atto680-labelled RNA was visualized with Odyssey Infrared Imager (Li-Cor Biosciences, Lincoln, NE, USA) and the RNA signals were quantified using ImageJ. The signals of unbound (A_59_), bound monomeric (A_59_:Nab2) and bound dimeric (A_59_:2×Nab2) complexes at each concentration of Nab2 were quantified to calculate the fractional saturation *Y*, as 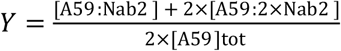, where [A_59_]_tot_ = 10 nM. This assumes that Nab2 has two binding sites on A_59_ RNA. The data points were fit in Origin to the Hill equation log 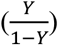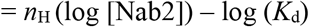, where *n*_H_ is the Hill coefficient and *K*_d_ is the dissociation constant. Note that the *K*_d_ derived from the *Y* intercept in a Hill plot is meaningless for interactions that do not occur with perfect cooperativity.

### Size exclusion chromatography multiangle light scattering (SEC-MALS)

Samples for SEC were mixed in polyadenylation buffer, incubated at 30 °C for 1 h, and centrifuged for 10 min at 10000 × g before loading to a Superdex 200 Increase 10/300 GL (bed volume 24 ml, GE Healthcare) column. The final concentrations of components in different samples were: A59 17 µM; A30 35 µM; Nab2-6xHis 31 µM; Nab2-6xHis + A59 (2:1) 15 µM + 7.5 µM; Nab2-6xHis + A59 (4:1) 16.5 µM + 4.125 µM; Nab2-6xHis + A30 15 µM + 15 µM. Total of 50-80 µg protein was loaded per sample in 45-75 µl volume. MALS analyses were performed on a MiniDAWN (Wyatt Technology) instrument with Optilab refractive index (RI) detector and Qasi-elastic light scattering (QELS) modules. The column and the HPLC detectors were placed in the cooling cabin with temperature ranging between 5 and 7 ºC. MALS (including QELS) and RI detectors were placed outside of the cabin and adjusted to 20 ºC. MALS analyses were done with Astra V 7.3.2 software (Wyatt Technology). The dn/dc and UV extinction coefficient values for the protein and RNA used as an input in Astra are reported in Supplemental Table 1. Molecular weights were calculated by using RI as concentration source. For conjugate analyses RI and UV were used as concentration sources.

### Formaldehyde crosslinking

RNA (0 or 100 nM) and Nab2-6xHis (250, 350 or 500 nM) were mixed in polyadenylation buffer using the Sarstedt Protein Low binding tubes and incubated for 1 h at 30 °C. 3% formaldehyde solution was then added at a final concentration of 0.3% (v/v) and incubated for 10 min at 30 °C. Cross-linking was stopped by adding 2 M glycine solution at a final concentration of 0.2 M and incubated for 5 min at 30 °C. 15 µl of the sample was transferred to a new tube and mixed with 10 µl of solution containing 2.5 × NuPAGE LDS sample buffer (Invitrogen) and 125 mM DTT. The mixture was incubated at 30 °C for 10 min, and 10 µl of sample was separated in NuPAGE 4-12% Bis-Tris gels (Invitrogen). The Atto680 fluorescence was subsequently visualized with Odyssey Infrared Imager (Li-Cor Biosciences, Lincoln, NE, USA), followed by silver staining of the gel using Thermo Scientific Pierce Silver Stain Kit according to the manufacturer’s instructions.

### Mass photometry

The reaction components, with concentrations detailed in the figure legends, were prepared in polyadenylation buffer using Sarstedt Protein Low binding tubes. The mixtures were incubated for 10 minutes at room temperature before recording 3-minute videos on a Refeyn TwoMP Auto mass photometer in manual mode with buffer-free focusing. Mass calibration was performed using a mixture containing 60 nM BSA (66 kDa) and 15 nM IgG (150, 300 kDa). Event histograms were analysed by Gaussian fitting using Refeyn Discover MP software.

### Yeast cultures and manipulations

Poly(A) tail length assays were conducted as reported in (Turtola et al. 2021). *MEX67-AA* strain (*tor1-1 fpr1::NAT RPL13-2xFKBP12*::*TRP1 MEX67-FRB*::*kanMX6 MATalpha*; Euroscarf) was transformed by the pESC-URA (pPgal) plasmids containing different *NAB2* constructs. Cells were cultured in synthetic complete (SC) medium lacking uracil and containing 2% raffinose and 0.1% glucose (SC-ura/raf). Three hours before heat shock 2% galactose was added to cell cultures which had an OD_600_ ∼0.5 to induce the protein expression from pESC-URA plasmids. Protein extracts were prepared from cells harvested 3 hours after adding the galactose. Cultures kept at 25°C were heat-shocked by rapidly mixing an equal volume of identical media pre-heated to 51°C and transferring the cultures to a 38 °C water bath. Rapamycin (final concentration 1 µg/ml; Life Technologies) was added to cultures 5 min before heat shock. Cells for RNA analysis were harvested by mixing equal volumes of culture and 96% ethanol pre-cooled on dry ice and centrifuging at 3000 × g for 3 min at 4 °C. The viability of *NAB2* mutations was evaluated by transforming the *nab2Δ::HIS3* p(*NAB2*/*CEN, URA3*) *ura3 leu2 trp1 MATalpha* strain (Marfatia et al. 2003) with p(*NAB2*/*CEN, LEU2*) plasmids. Plasmid transformations were performed using lithium acetate/PEG method with denatured salmon sperm DNA as a carrier. To conduct the growth assays, transformed yeast cells were grown over night in SC medium containing 2 % glucose and lacking leucine (SC-leu/glu) or in SC medium containing glucose and lacking uracil and leucine (SC-ura-leu/glu). Cell densities were then measured and adjusted to OD_600_=1, and 10-fold serial dilutions were spotted on the SC-leu/glu (or SC-ura-leu/glu) and SC/glu 5-FOA (1 mg/ml) agar plates, and incubated at the indicated temperatures before imaging at the indicated times. The transformed cells were cured from p(*NAB2*/*CEN, URA3*) plasmids by two passages on SC/glu 5-FOA plates, resulting in strains with a genotype *nab2Δ::HIS3* p(*NAB2*/*CEN, LEU2*) *ura3 leu2 trp1 MATalpha*, with *NAB2* genes bearing the indicated mutations. The Nab2 protein levels were determined from these strains that were cultured at 30 °C in SC/glu medium to an OD_600_ of approximately 0.6.

### RNA extraction

RNA extraction was performed with the hot phenol method. Briefly, a frozen cell pellet was resuspended in TES-buffer (1% SDS, 5 mM EDTA, 10 mM Tris-HCl, pH 7.5) and an equal volume of 0.1 M citrate buffered phenol pH 4.3 (Sigma P4682). The mix was incubated at 65 °C for 40 min under 1400 rpm shaking, followed by centrifugation at 16000 × g for 10 min at 4 °C. The aqueous phase was extracted again with acidic phenol at 65 °C for 20 min, and twice with chloroform at room temperature. RNA was precipitated with ethanol (final 70%) and 60 mM LiCl over-night. The pellet was washed with EtOH and resuspended in water.

### RNaseH Northern Blotting

RNaseH northern blotting was performed as reported in (Turtola et al. 2021). Briefly, 20 μg of total RNA was combined with 2 μM DNA oligonucleotide (DL163 (Libri et al. 2002); oligonucleotide sequences are listed in **Supplemental Table S4**) complementary to the *HSP104* 3′ region in annealing buffer (50 mM Tris-HCl, pH 8.3, 50 mM KCl) in a total volume of 12 μl. The mix was incubated at 85 °C for 2 min and slowly cooled to 37 °C. Subsequently, annealing reactions were supplemented with 8 μl mix pre-heated to 37 °C and containing 2.5 U of RNaseH (New England Biolabs), 2.5 × RNaseH reaction buffer (1× buffer: 50 mM Tris-HCl, 75 mM KCl, 3 mM MgCl_2_, 10 mM DTT, pH 8.3), 25 mM DTT and 4 U RiboLock RNase inhibitor (Thermo Scientific). The mix was incubated at 37 °C for 30 min, followed by addition of 100 μl absolute ethanol and 20 μl of solution containing 600 mM sodium acetate (pH 5.3), 10 mM EDTA, 5 μg tRNA (Roche 28473522) and 5 μg glycogen. RNA was then precipitated at -20 °C. The pellet was washed with 70% ethanol and resuspended in RNA loading buffer (formamide, 10 mM Tris-HCl pH 8.0, 5 mM EDTA, 0.02% xylene cyanol). Samples were separated on 6% urea-polyacrylamide gels by electrophoresis, transferred to Hybond-N+ membrane (GE Healthcare RPN203B), hybridized over-night at 50 °C with 5′ terminally ^32^P-labelled DNA oligonucleotide probes DL164 (Libri et al. 2002); and MS618 (Schmid et al. 2015) in ULTRA-Hyb Oligo Hybridization buffer (Invitrogen AM8663), washed 4 times with 2 × SSC (300 mM NaCl, 30 mM trisodium citrate, pH 7.0) containing 0.5% SDS, each time rotating for 30 min at 42 °C, and exposed to phosphorimager screen. Images were processed and quantitated with ImageJ software. The indicated poly(A) tail lengths were approximated from a DNA size marker.

### Western blotting

Cells equivalent to 5 OD units were lysed by vortexing with glass beads in 8 M urea for 5 minutes at 4 °C, followed by incubation at 95 °C for 10 minutes. Lysate was clarified by centrifugation. Total protein concentration was determined by a Bradford assay, and equal amounts protein were loaded onto an SDS-PAGE gel. Western blotting was carried out with standard procedures using antibodies for Nab2 (HL831 3F2; (Anderson et al. 1993)), Rpb3 (1Y26; Abcam), and FlagM2 (F1804; Sigma-Aldrich), which were detected using either HRP-conjugated secondary anti-mouse polyclonal goat immunoglobulins (Dako) or Alexa Fluor 680-conjugated secondary anti-mouse cross-adsorbed secondary antibody (A21057; Invitrogen).

## Supporting information

Gabs et al_Supplemental figures and tables

## COMPETING INTEREST STATEMENT

The authors declare no competing interests.

## ACKNOWLEDGEMENTS

We thank Anita Corbett for the *nab2Δ* p(*NAB2*/*CEN, URA3*) strain and Françoise Stutz for providing the pAC1039 plasmid. Dorthe Caroline Riishøj and Andrei Zupnik are acknowledged for expert technical assistance, and Georgiy Belogurov’s lab for sharing essential equipment. Kalle Sipilä, Søren Lykke-Andersen and the members of T.H. J., L.A.P. and M.T. labs are thanked for helpful discussions. Kristian Koski is thanked for technical help in SEC-MALS experiments and analyses. The use of the facilities and expertise of the Biocenter Oulu Structural Biology core facility, a member of Biocenter Finland, Instruct-ERIC Centre Finland and FINStruct, is gratefully acknowledged. Financial support was provided by Instruct-ERIC (PID: 26089). Turku Protein Core is acknowledged for providing access to mass photometer and kinetic instruments. The facilities and expertise of the Structural Bioinformatics Laboratory, Åbo Akademi University, a member of Turku Protein Core, FINStruct, and Biocenter Finland are gratefully acknowledged. This work was supported by Research Council of Finland (grants 349698 and 353682; M.T.), Magnus Ehrnrooth foundation (M.T.), the Medical Research Council (MRC), as part of United Kingdom Research and Innovation (MRC file reference number MC_U105192715; L.A.P.), and the Wellcome Trust (Grant reference number 225217/Z/22/Z; L.A.P.).

For the purpose of open access, the MRC Laboratory of Molecular Biology has applied a CC BY public copyright licence to any Author Accepted Manuscript version arising from this submission.

## AUTHOR CONTRIBUTIONS

Conceptualization, M.T., L.A.P. and S.H.M.; Methodology, M.T. and S.H.M.; Formal Analysis, M.T., S.H.M. and E.A.S.; Investigation, E.G., M.T., S.H.M., E.A.S., A.V.; Resources, M.T., L.A.P., T.H.J., A.M.M.; Writing – Original Draft, M.T.; Writing – Review & Editing, T.H.J., S.H.M., L.A.P. and A.M.M.; Visualization, M.T. and S.H.M.; Supervision, M.T.; Funding Acquisition, M.T., T.H.J., L.A.P.

## Notes

### Competing Interest Statement

The authors have declared no competing interest.

