## Supplementary material for "A kinetic ruler controls mRNA poly(A) tail length": Gabs et al_Supplemental figures and tables

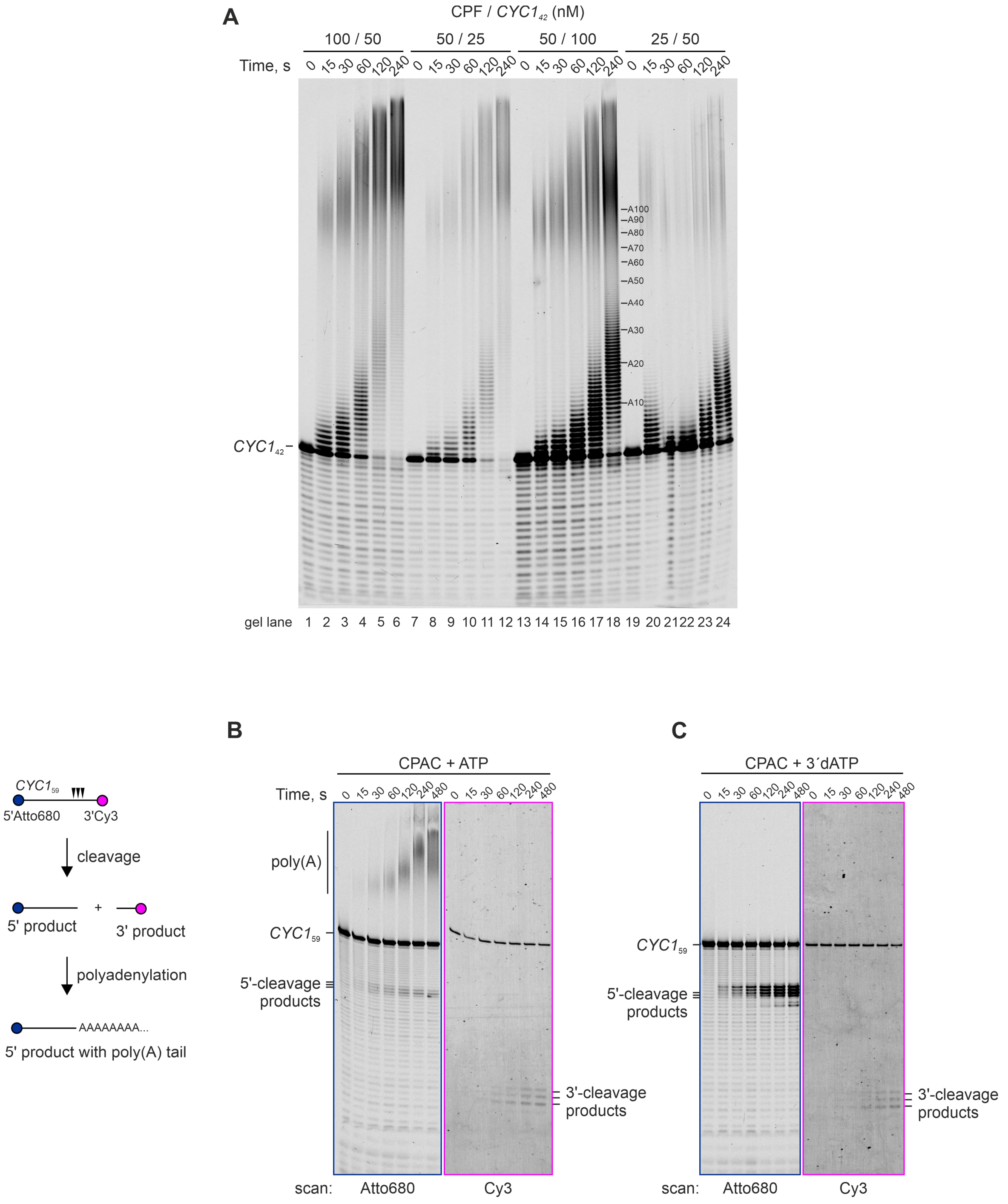


**Supplemental Figure S1.** (*A*) The effect of RNA and CPF concentrations on polyadenylation. The concentrations of CF IA (500 nM), CF IB (500 nM) and ATP (2 mM) were the same in all reactions. (*B*) Cleavage and polyadenylation. A dual-labelled *CYC1*$_{59}$ RNA substrate with 5′Atto680 and 3′Cy3 dyes was used for tracking the 5′ and 3′ products. The RNA was pre-incubated with CF IA and CF IB, before the start of the reactions by the simultaneous addition of CPF and ATP. The scans of the same gel area are shown for Atto680 (*left*; blue outline) and Cy3 (right; magenta outline) channels. (*C*) As in *B* but 3′dATP was included in the reactions to prevent polyadenylation.

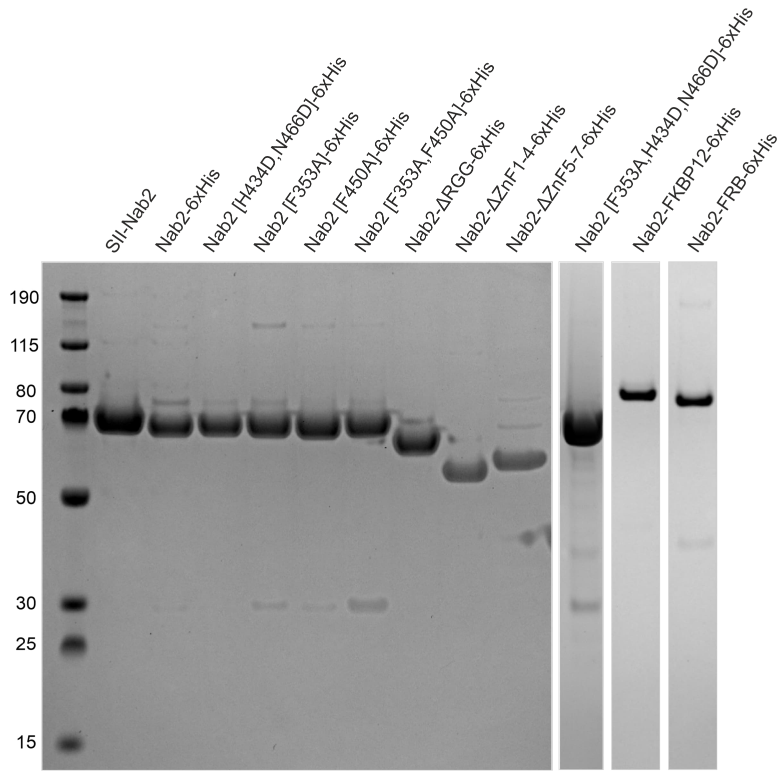


**Supplemental Figure S2.** Purified Nab2 proteins used in Figures 2-6 separated by SDS-PAGE and visualised by Coomassie staining. SII-Nab2 was expressed in *Sf9* insect cells. Other proteins were expressed in *E. coli*.

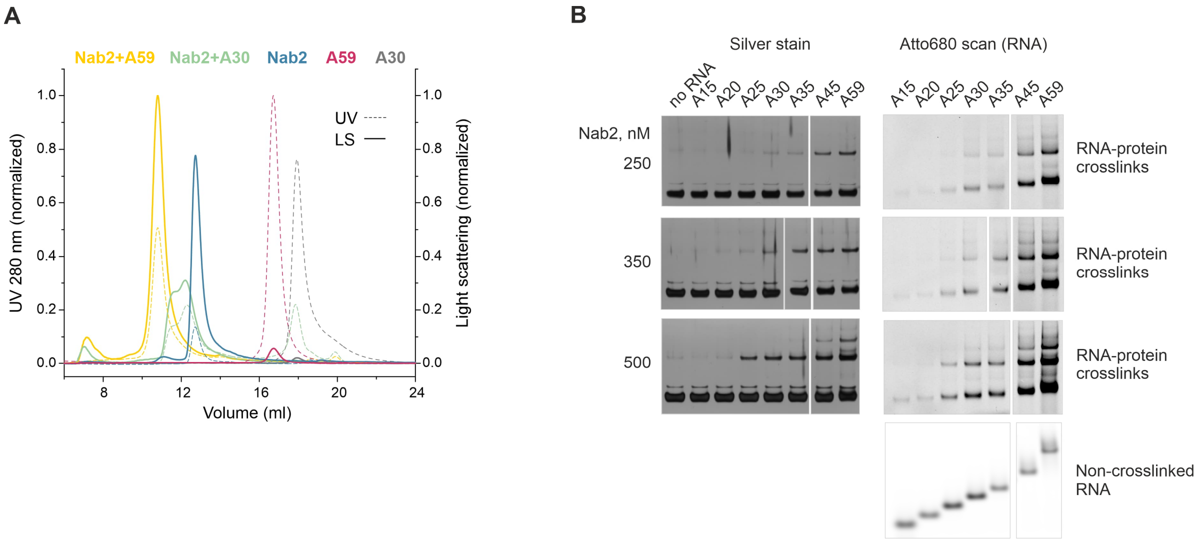


**Supplemental Figure S3.** (*A*) UV (280 nm) and light scattering traces of the SEC-MALS analyses for $A_{59}$ (magenta), $A_{30}$ (light grey), Nab2 (blue) and Nab2 mixed with either $A_{59}$ (yellow) or $A_{30}$ (green). Light scattering intensities are displayed (y-axis on the right side) along the elution volume (x-axis). The absorbance is displayed by dashed lines with corresponding colours (y-axis on the left side). All absorbance values were normalized to the highest value in the $A_{59}$ RNA sample. All light scattering values were normalized to the highest value in the Nab2+$A_{59}$ RNA sample. (*B*) Formaldehyde crosslinking of Nab2 (250, 350, 500 nM) with Atto680-labelled poly(A) RNAs (100 nM) of varying lengths. The panels were cropped together from two gels exposed at the same time.

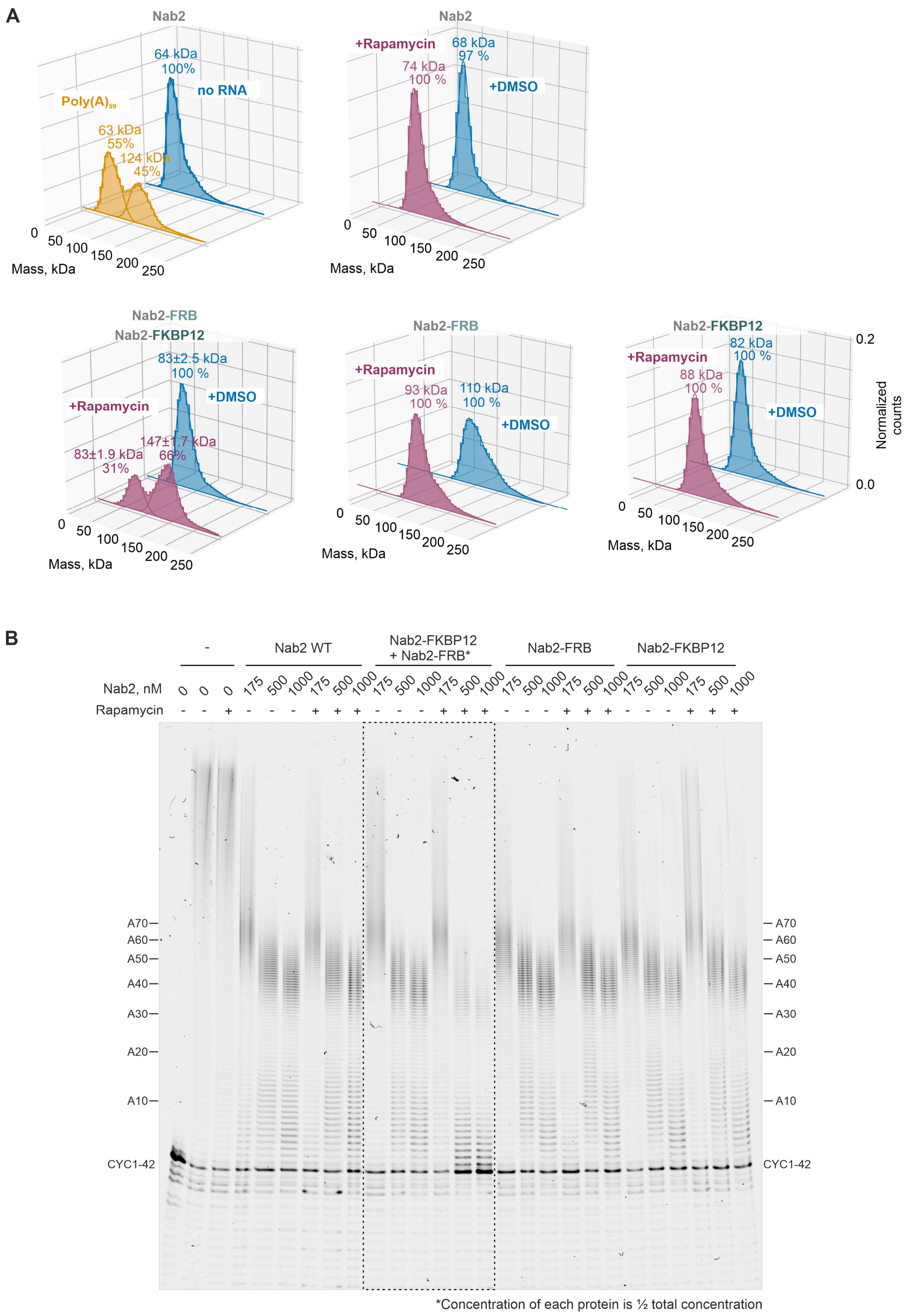


**Supplemental Figure S4.** (*A*) Mass photometric analysis of the effects of DMSO and rapamycin (1 µM) for the oligomerization state of Nab2, Nab2-FKBP12 and Nab2-FRB. The total concentration of Nab2 in all experiments was 250 nM. The combination of Nab2-FKBP12 and Nab2-FRB (each 125 nM) is the same as shown in Fig. 4B. Nab2 (250 nM) with $A_{59}$ (50 nM) on the left is shown as a positive control for RNA-dependent dimerization of Nab2. The gaussian fits of the count-normalized histograms are shown as lines with the peak positions (in kDa) and the percentage of total peak counts displayed above. (*B*) The effects of rapamycin and the Nab2-FRB and Nab2-FKBP12 fusion proteins (added individually or together) on *CYC1*$_{42}$ polyadenylation by the CPAC. Nab2 proteins were incubated with DMSO (-rapamycin) or with 1 µM rapamycin before being added to the polyadenylation reactions together with ATP. The reactions were stopped after 4 minutes. The marked gel area is the same as displayed in Fig. 4C.

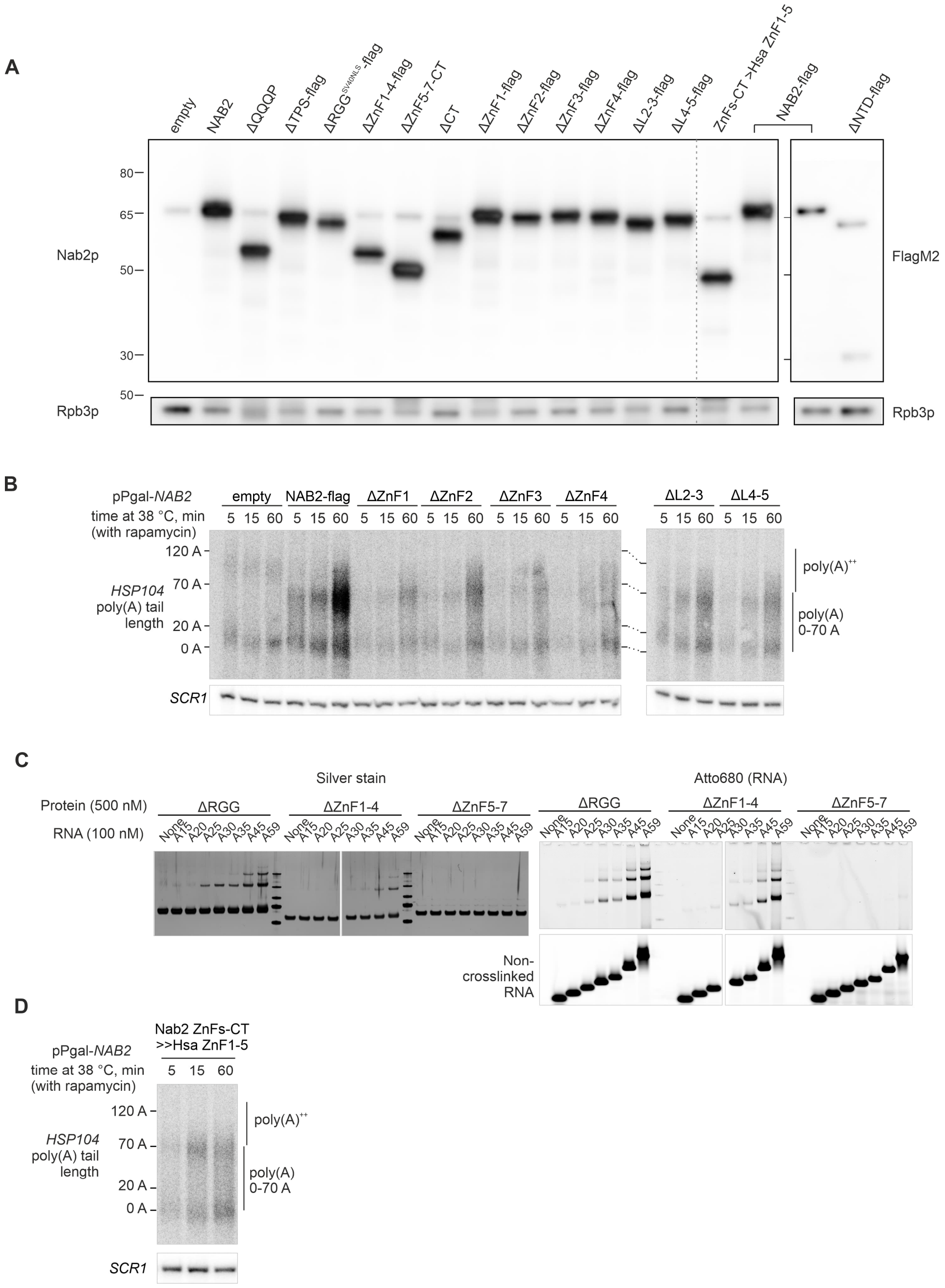


**Supplemental Figure S5.** (*A*) Western blot analysis of the Nab2 and Rpb3 protein levels in protein extracts from *MEX67-AA* cells over-expressing different Nab2 mutant variants used in Fig. 5B, Supplemental Fig. 5B and Supplemental Fig. 5D. The upper left panels were probed with an anti-Nab2 antibody (Anderson et al. 1993) that binds to the NTD. The ΔNTD-flag variant, and for comparison the wild-type Nab2-flag, were detected with an anti-FlagM2 antibody (upper right panel). Rpb3 was detected afterwards from the same membranes that were cut at the 50 kDa marker. Note that the endogenously expressed Nab2 protein is visible as a faint band migrating at the level of the 65 kDa marker. (*B*) Nuclear *HSP104* mRNA poly(A) tail lengths in cells expressing additional mutant variants of Nab2. See Fig. 5A-B for experimental details. (*C*) Additional data for Fig. 5D showing the silver stained and Atto680 scans of formaldehyde crosslinked Nab2 truncation mutants (500 nM) incubated with Atto680-labelled poly(A) RNAs (100 nM) of varying lengths. The panels were cropped together from two gels that were simultaneously exposed. (*D*) Nuclear *HSP104* mRNA poly(A) tail lengths in cells expressing a chimeric protein where the ZnF1-7-CT region of *S. cerevisiae* Nab2 is replaced by ZnF1-5 of human ZC3H14.

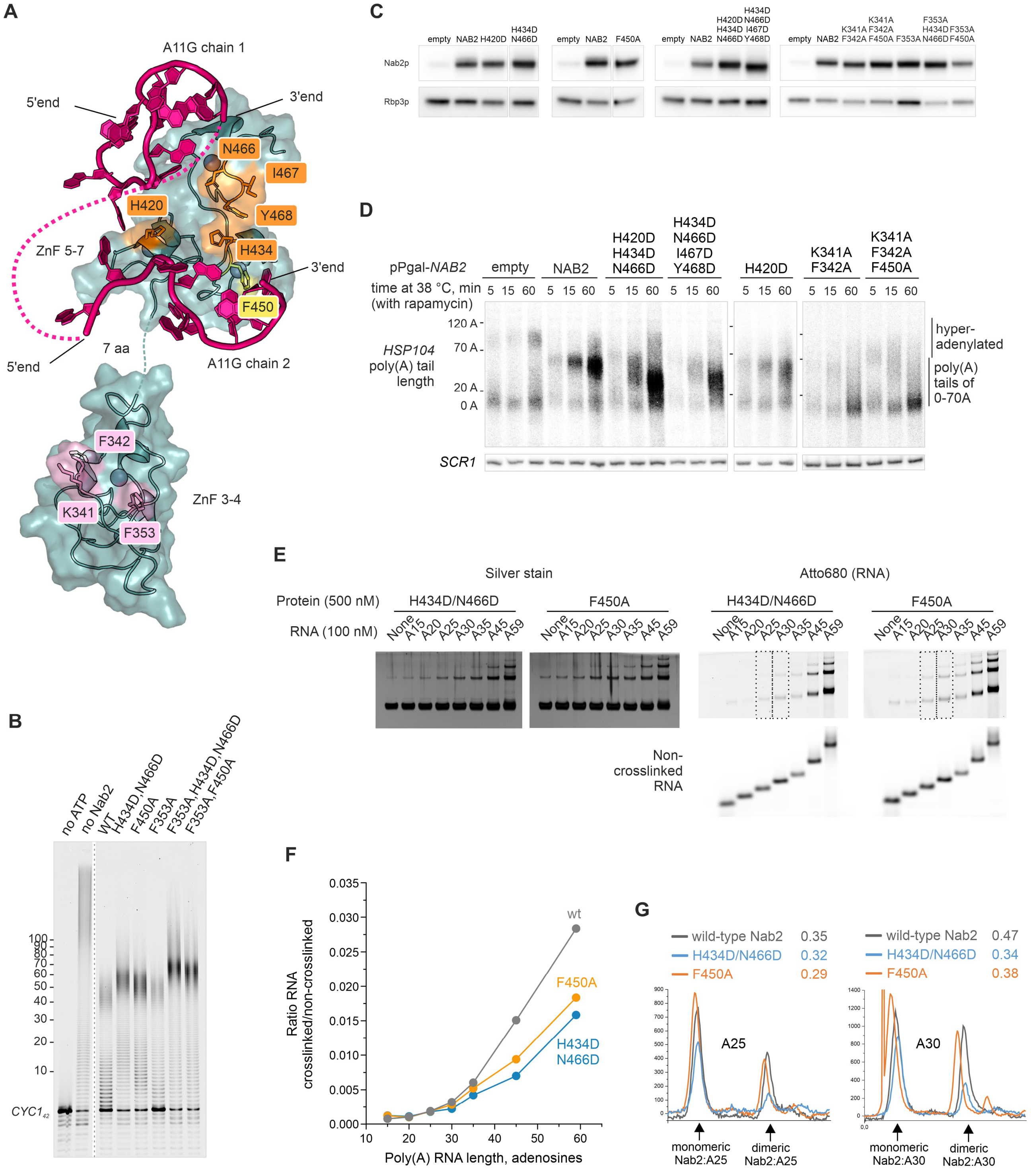


**Supplemental Figure S6.** (*A*) A composite model of Nab2 showing one ZnF5-7 domain bound to two chains of $A_{11}$G RNA (*top*, PDB 5L2L; (Aibara et al. 2017)) and one ZnF3-4 domain (*bottom*, PDB 3ZJ2; (Martínez-Lumbreras et al. 2013)). The second ZnF5-7 domain is removed in order to display the dimeric interface surface. The protein residues mutated in this study are shown as sticks. The mutated dimer interface residues are highlighted in orange and the mutated residues interacting directly with RNA are highlighted in ZnF6 in yellow (F450) or in ZnF3 in pink (K341, F342, F353). The 5′ and 3′ ends of the RNAs are indicated. The dotted magenta line depicts a hypothetical path to connect two ends by an intervening RNA chain. (*B*) The effect of purified Nab2 point mutants (1000 nM) on CPAC-mediated *CYC1*$_{42}$ polyadenylation. (*C*) Western blot analysis of the Nab2 and Rpb3 protein levels in protein extracts from *MEX67-AA* cells over-expressing different Nab2 mutant variants used in Fig. 6C and Supplemental Fig. 6D. Gel panels with narrow spacing were cropped from the same gels. (*D*) Nuclear *HSP104* mRNA poly(A) tail lengths in cells expressing mutant variants of Nab2. See Fig. 5A-B for experimental details. (*E*) formaldehyde crosslinked Nab2 point mutants (500 nM) incubated with Atto680-labelled poly(A) RNAs (100 nM) of varying lengths. The signal intensity scans from the dashed areas are shown in *G*. (*F*) Quantification of poly(A) RNA-Nab2 crosslinks from *E*. The wild-type Nab2 (500 nM) data presented in Fig. 3E is replicated here. (*G*) Signal intensity scans of H434D/N466D and F450A mutants crosslinked with A25 and A30 RNAs. The corresponding scans of wild-type Nab2 crosslinks are shown for comparison. The intensities of the crosslinked RNAs were normalized to the total intensities of the non-crosslinked RNAs. The ratios of multimeric vs. total RNA-protein crosslinks are shown on top.

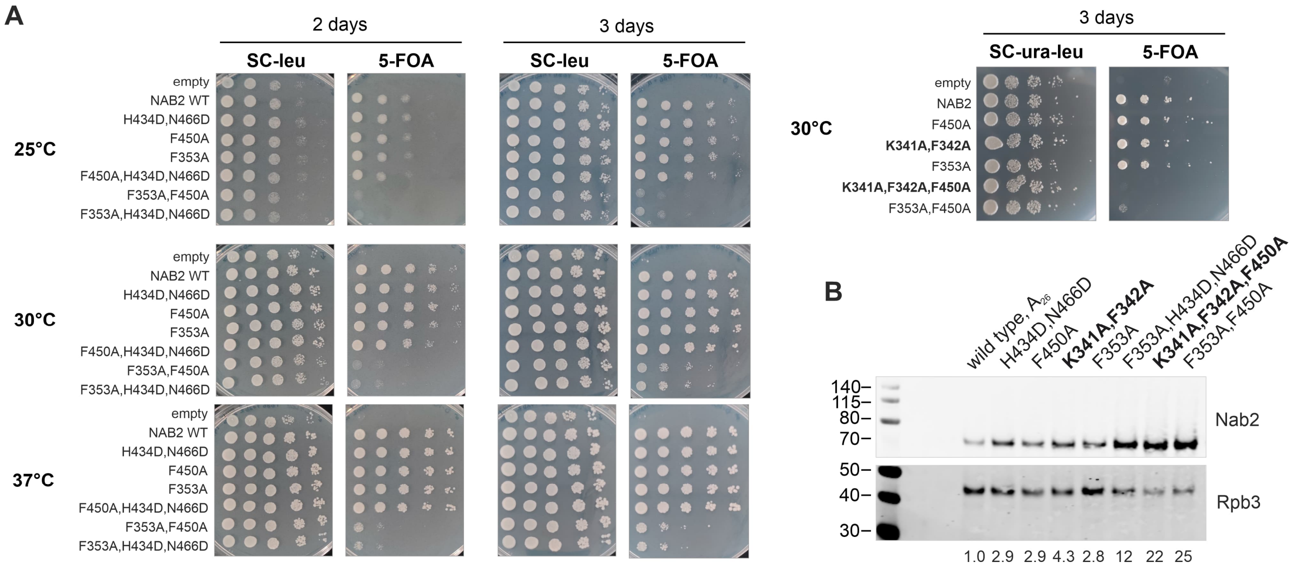


**Supplemental Figure S7.** (*A*) Growth assays for the *Δnab2::HIS3* pURA3/*NAB2* strain transformed by the p(*NAB2*/*CEN, LEU2*) plasmids carrying mutations in *NAB2*. The cells grown in SC medium lacking leucine (SC-leu), or uracil and leucine (SC-ura-leu) were adjusted to $OD_{600}$=1, 10-fold serial dilutions were spotted on the SC-leu (or SC-ura-leu) and SC 5-FOA agar plates incubated at 25, 30 or 37 °C, and imaged at the indicated times. (*B*) Western blot analysis of the Nab2 and Rpb3 protein levels in the *Δnab2::HIS3* p(*NAB2*/*CEN, LEU2*) strains bearing the indicated mutations in *NAB2*. The mean of Nab2/Rpb3-ratios from two experiments normalized to the wild-type is shown below.

### SUPPLEMENTAL TABLES

#### Supplemental Table S1. Size exclusion chromatography-multiangle light scattering data

| **Protein** | Nab2-6xHis* | Nab2-6xHis* | Nab2-6xHis* | Nab2-6xHis* | - | - |
|---|---|---|---|---|---|---|
| **RNA** | A59** | A59** | A30*** | - | A59** | A30*** |
| Molar ratio (protein:RNA) in the sample | 2:1 | 4:1 | 2:2 | n/a | n/a | n/a |
| **MW based on refractive index (RI)** | | | | | | |
| MW (kDa) | 135.3 | 143.9 | 82.2 | 59.4 | 19.2 | 11.2 |
| **Conjugate analysis, MW based on RI and UV extinction coefficient** | | | | | | |
| MW (kDa) | 135.6 | 144 | 83.1 | n/a | n/a | n/a |
| MW Protein (kDa) | 119.3 | 128 | 68.9 | n/a | n/a | n/a |
| MW Modifier (kDa) | 16.3 | 15.7 | 14.3 | n/a | n/a | n/a |

*Mr 59.143 kDa, dn/dc [ml/g] 0.185, UVext [ml/(mg×cm)] 0.224
**Mr 19.441 kDa, dn/dc [ml/g] 0.18, UVext [ml/(mg×cm)] 11.6
***Mr 9.894 kDa, dn/dc [ml/g] 0.18, UVext [ml/(mg×cm)] 5.90

#### Supplemental Table S2. Plasmids

| **Identifier** | **Original name** | **Description** | **Source** |
|---|---|---|---|
| **Baculovirus integration vectors** | | | |
| CPF polymerase module | P20-3 | pBIG1a (CFT1 N997D, PFS2-SII, YTH1, FIP1, PAP1) | Hill et al. 2019 |
| CPF nuclease module | P20-6 | pBIG1b (CFT2, YSH1, MPE1) | Kumar et al. 2021 |
| CPF phosphatase module | P27-37, clone AC2 | pBIG2ab (SSU72, PTI1, GLC7, REF2-SII, SWD2, PTA1) | Kumar et al. 2021 |
| CF IA | P20-24 | pBIG1c (8xHisRNA14, RNA15, SII-PCF11, CLP1) | Kumar et al. 2021 |
| Nab2 | P21-63 | pACEBac1 (SII-NAB2) | Turtola et al. 2021 |
| ***E. coli* expression plasmids** | | | |
| Nab2 | pEG005 | pET28b C-terminal 6xHis | This study |
| Nab2-FKBP12 | pEG025 | pET28b C-terminal FKBP12 (H. sapiens FKBP12 residues 1-107), C-terminal 6xHis | This study |
| Nab2-FRB | pEG026 | pET28b C-terminal FRB (H. sapiens mTOR residues 2021-2113, T2098L), C-terminal 6xHis | This study |
| Nab2_ΔRGG | pEG007 | pET28b [201-255 deleted], C-terminal 6xHis | This study |
| Nab2_ΔZnF1-4 | pEG006 | pET28b [262-389 deleted], C-terminal 6xHis | This study |
| Nab2_ΔZnF5-7 | pEG012 | pET28b [390-485 deleted], C-terminal 6xHis | This study |
| Nab2_N466D,H434D | pEG008 | pET28b N466D,H434D, C-terminal 6xHis | This study |
| Nab2_F450A | pEG009 | pET28b F450A, C-terminal 6xHis | This study |
| Nab2_F353A | pEG010 | pET28b F353A, C-terminal 6xHis | This study |
| Nab2_F353A,H434D,N466D | pEG017 | pET28b F353A,H434D,N466D, C-terminal 6xHis | This study |
| Nab2_F353A,F450A | pEG011 | pET28b F353A,F450A, C-terminal 6xHis | This study |
| CF IB | P2-43 | pOPINB 6xHis-HRP1 | Casañal et al. 2017 |
| ***NAB2* in pPgal plasmid*** | | | |
| empty | pESC-URA | empty plasmid | Stratagene |
| NAB2 | p429 | wild type NAB2 | Tudek et al., 2018 |
| ΔNTD | p437 | [4-97 deleted], C-terminal flag-tag | This study |
| ΔQQQP | p433 | [104-169 deleted] | This study |
| ΔTPS | p441 | [170-200 deleted], C-terminal flag-tag | This study |
| ΔRGG | p442 | [201-255 deleted]; SV40 NLS-GlyGly inserted at the N-terminus, C-terminal flag-tag | This study |
| ΔZnF1-4 | p440 | [262-389 deleted], C-terminal flag-tag | This study |
| ΔZnF5-7-CT | p432 | [395-525 deleted] | This study |
| ΔCT | p450 | [486-525 deleted] | This study |
| ΔZnF1 | p443 | [262-278 deleted], C-terminal flag-tag | This study |

| | | | |
|---|---|---|---|
| ΔZnF2 | p444 | [283-300 deleted], C-terminal flag-tag | This study |
| ΔZnF3 | p446 | [340-355 deleted], C-terminal flag-tag | This study |
| ΔZnF4 | p447 | [371-386 deleted], C-terminal flag-tag | This study |
| ΔL2-3 | p445 | [305-333 deleted], C-terminal flag-tag | This study |
| ΔL4-5 | p448 | [391-408 deleted], C-terminal flag-tag | This study |
| NAB2-flag (wild type) | p434 | C-terminal flag-tag | This study |
| H434D, N466D | p482 | H434D,N466D | This study |
| F450A | p495 | F450A | This study |
| F353A | p517 | F353A | This study |
| F353A, H434D, N466D | p518 | F353A,H434D,N466D | This study |
| F353A, F450A | p508 | F353A,F450A | This study |
| K341A, F342A | p516 | K341A,F342A | This study |
| K341A, F342A, F450A | p507 | K341A,F342A,F450A | This study |
| H420D | p478 | H420D | This study |
| H420D, H434D, N466D | p496 | H420D,H434D,N466D | This study |
| H434D, N466D, I467D, Y468D | p497 | H434D,N466D,I467D,Y468D | This study |
| ZnF1-7-CT >> Hsa ZnF1-5 | p458 | SceNab2 amino acids 1-255 fused to HsaZC3H14 amino acids 594-736 | This study |
| **p(*NAB2/CEN, LEU2*) plasmid**** | | | |
| empty | p513H | p(*CEN, LEU2*); empty plasmid | This study |
| NAB2 | p519 | wild type NAB2 | This study |
| N466D,H434D | p525 | N466D,H434D | This study |
| F450A | p528 | F450A | This study |
| F353A | p530 | F353A | This study |
| K341A,F342A | p529 | K341A,F342A | This study |
| N466D,H434D,F353A | p537 | N466D,H434D,F353A | This study |
| F450A,F353A | p535 | F450A,F353A | This study |
| F450A,K341A,F342A | p534 | F450A,K341A,F342A | This study |
| NAB2 (wild type) | pMT105 | A26>A11 (autoregulatory downstream sequence) | This study |
| NAB2 (wild type) | pMT106 | A26>A16 (autoregulatory downstream sequence) | This study |
| NAB2 (wild type) | pAV002 | A26>A21 (autoregulatory downstream sequence, spontaneous mutation in p513 clone F) | This study |
| NAB2 (wild type) | pMT107 | A26>A33 (autoregulatory downstream sequence) | This study |
| NAB2 (wild type) | pMT108 | A26>A42 (autoregulatory downstream sequence) | This study |
| **pESC-URA 2µ, ori(f1), ori(pUC), URA3 , Ampr, TCYC1, MCS, PGAL1, PGAL10, MCS, TADH1; NAB2-genes cloned under PGAL10 with EcoRI and NotI* | | | |
| ***pRSII415 LEU2, CEN6/ARSH4, pBluescript II SK+ NAB2 promoter (400 bp)_ EcoRI-NotI_NAB2 downstream sequence (400 bp); NAB2 genes cloned with EcoRI and NotI* | | | |

### Supplemental Table S3. Primers

| | Plasmid number | Sequence [deletions], point mutations, modifications | Two-step PCR | PCR template | Primer sequence (5'-3') | Primer Description |
|---|---|---|---|---|---|---|
| ***NAB2* in pPgal plasmid*** | | | | | | |
| ΔNTD | p437 | [4-97], C-terminal flag-tag | n.a. | p429 | ACAGCAGAATTCATGTCTCAAAGCTTGGGACAATCGGATATC | Forw_EcoRI-[MSQ-Nab2 delta NTD 4-97] |
| | | | | | ACAAACGCGGCCGCGTTCATTTCCGTATCTTGTTCTTG | Rev NotI-[C-term of Nab2], no term codon |
| ΔQQQP | p433 | [104-169] | n.a. | pAC1039 | gtccactgGAATTCATGTCTCAAGAACAGTACACAGAAAA | Fwd for Nab2 ORF amplification with EcoRI site |
| | | | | | gctgacctGCGGCCGCTCAGTTCATTTCCGTATCTTGTTC | Rev for Nab2 ORF amplification with NotI site |
| ΔTPS | p441 | [170-200], C-terminal flag-tag | first step, N | p429 | ATGTCTCAAGAACAGTACACAG | Forw Nab2 N-term |
| | | | | | GGCTGTTATCATTCTGGGTCCCTAGTTGT | delTPS (rev A) |
| | | | first step, C | p429 | GACCCAGAATGATAACAGCCAAAGGTTCAC | delTPS (fw B) |
| | | | | | GTTCATTTCCGTATCTTGTTCTTG | Rev Nab2 C-term |
| | | | second step | N+C (from first step) | gtccactgGAATTCATGTCTCAAGAACAGTACACAGAAAA | Fwd for Nab2 ORF amplification with EcoRI site |
| | | | | | ACAAACGCGGCCGCGTTCATTTCCGTATCTTGTTCTTG | Rev NotI-[C-term of Nab2], no term codon, can be cloned in frame with FLAG in pESC |
| ΔRGG | p442 | [201-255]; SV40 NLS-GlyGly inserted at the N-terminus, C-terminal flag-tag | first step, N | p429 | ATGTCTCAAGAACAGTACACAG | Forw Nab2 N-term |
| | | | | | CTTTCTTGGTTACAGGCGCAAACTGAGG | delRGG-ver2 (rev A) |
| | | | first step, C | p429 | TGCGCCTGTAACCAAGAAAGAGGGGCGT | delRGG-ver2 (fw B) |
| | | | | | GTTCATTTCCGTATCTTGTTCTTG | Rev Nab2 C-term |
| | | | second step | N+C (from first step) | ACAGCAGAATTCATGCCAAAAAAAAAAAGAAAAGTTGGCGGCTCTCAAGAACAGTACACAGAAAAC | Forw_EcoRI-[N-term of SV40*NLS-GG-Nab2 N-term] |
| | | | | | ACAAACGCGGCCGCGTTCATTTCCGTATCTTGTTCTTG | Rev NotI-[C-term of Nab2], no term codon, can be cloned in frame with FLAG in pESC |
| ΔZnF1-4 | p440 | [262-389], C-terminal flag-tag | first step, N | p429 | ATGTCTCAAGAACAGTACACAG | Forw Nab2 N-term |
| | | | | | TGATCTTCGAACGCCCCTCTTTCTTGGTTG | Rev Nab2 residue ...261+10 bp overhang complementary to residues 390… -->2-step PCR deletion of ZnF1-4 |
| | | | first step, C | p429 | AGAGGGGCGTTCGAAGATCAAGGAAGTAAAACC | Forw Nab2 residue 390…+10 bp overhang complementary to residues …261 -->2-step PCR deletion of ZnF1-4 |
| | | | | | GTTCATTTCCGTATCTTGTTCTTG | Rev Nab2 C-term |
| | | | second step | N+C (from first step) | gtccactgGAATTCATGTCTCAAGAACAGTACACAGAAAA | Fwd for Nab2 ORF amplification with EcoRI site |
| | | | | | ACAAACGCGGCCGCGTTCATTTCCGTATCTTGTTCTTG | Rev NotI-[C-term of Nab2], no term codon, can be cloned in frame with FLAG in pESC |
| ΔZnF5-7-CT | p432 | [395-525] | n.a. | p429 | gtccactgGAATTCATGTCTCAAGAACAGTACACAGAAAA | Fwd for Nab2 ORF amplification with EcoRI site |
| | | | | | ACAACAGCGGCCGCTCACAATGAAGAGTGGGCCTTTC | Rev_Truncation of Nab2 ZnF 5-7, residues 390-525. Introduce termination codon and NotI site |
| ΔCT | p450 | [486-525] | n.a. | p429 | gtccactgGAATTCATGTCTCAAGAACAGTACACAGAAAA | Fwd for Nab2 ORF amplification with EcoRI site |
| | | | | | ACAAACGCGGCCGCTCAAGCGCCTTTCTTTTCCG | delCT[486-525]-NotI-without flag_rev |
| ΔZnF1 | p443 | [262-278], C-terminal flag-tag | first step, N | p429 | ATGTCTCAAGAACAGTACACAG | Forw Nab2 N-term |
| | | | | | CCTTAGTTGGACGCCCCTCTTTCTTGGT | delZnF1 (rev A) |
| | | | first step, C | p429 | AGAGGGGCGTCCAACTAAGGTATGTAATGAATATCC | delZnF1 (fw B) |
| | | | | | GTTCATTTCCGTATCTTGTTCTTG | Rev Nab2 C-term |
| | | | second step | N+C (from first step) | gtccactgGAATTCATGTCTCAAGAACAGTACACAGAAAA | Fwd for Nab2 ORF amplification with EcoRI site |
| | | | | | ACAAACGCGGCCGCGTTCATTTCCGTATCTTGTTCTTG | Rev NotI-[C-term of Nab2], no term codon, can be cloned in frame with FLAG in pESC |
| ΔZnF2 | p444 | [283-300], C-terminal flag-tag | first step, N | p429 | ATGTCTCAAGAACAGTACACAG | Forw Nab2 N-term |
| | | | | | CTTCATTTGGTACCTTAGTTGGGTGTGCATG | delZnF2 (rev A) |
| | | | first step, C | p429 | CCCAACTAAGGTACCAAATGAAGATGAAGAGTTGATG | delZnF2 (fw B) |
| | | | | | GTTCATTTCCGTATCTTGTTCTTG | Rev Nab2 C-term |
| | | | second step | N+C (from first step) | gtccactgGAATTCATGTCTCAAGAACAGTACACAGAAAA | Fwd for Nab2 ORF amplification with EcoRI site |
| | | | | | ACAAACGCGGCCGCGTTCATTTCCGTATCTTGTTCTTG | Rev NotI-[C-term of Nab2], no term codon, can be cloned in frame with FLAG in pESC |
| ΔZnF3 | p446 | [340-355], C-terminal flag-tag | first step, N | p429 | ATGTCTCAAGAACAGTACACAG | Forw Nab2 N-term |
| | | | | | CTGGTGTTGGCAGAACGATACCAGTTTGTACC | delZnF3 (rev A) |
| | | | first step, C | p429 | TATCGTTCTGCCAACACCAGCAAATGAAG | delZnF3 (fw B) |
| | | | | | GTTCATTTCCGTATCTTGTTCTTG | Rev Nab2 C-term |
| | | | second step | N+C (from first step) | gtccactgGAATTCATGTCTCAAGAACAGTACACAGAAAA | Fwd for Nab2 ORF amplification with EcoRI site |

| | | | | | | |
|---|---|---|---|---|---|---|
| | | | | | ACAAACGCGGCCGCGTTCATTTCCGTATCTTGTTCTTG | Rev NotI-[C-term of Nab2], no term codon, can be cloned in frame with FLAG in pESC |
| ΔZnF4 | p447 | [371-386], C-terminal flag-tag | first step, N | p429 | ATGTCTCAAGAACAGTACACAG | Forw Nab2 N-term |
| | | | | | ACAATGAAGACCACATTAGATCAATGACTTTCG | delZnF4 (rev A) |
| | | | first step, C | p429 | ATTGATCTAATGTGGTCTTCATTGTCGAAGATCAAGG | delZnF4 (fw B) |
| | | | | | GTTCATTTCCGTATCTTGTTCTTG | Rev Nab2 C-term |
| | | | second step | N+C (from first step) | gtccactgGAATTCATGTCTCAAGAACAGTACACAGAAAA | Fwd for Nab2 ORF amplification with EcoRI site |
| | | | | | ACAAACGCGGCCGCGTTCATTTCCGTATCTTGTTCTTG | Rev NotI-[C-term of Nab2], no term codon, can be cloned in frame with FLAG in pESC |
| ΔL2-3 | p445 | [305-333], C-terminal flag-tag | first step, N | p429 | ATGTCTCAAGAACAGTACACAG | Forw Nab2 N-term |
| | | | | | CAGTTTGTACATCTTCATTTGGATGTAAAACTCAC | delLinker2-3 (rev A) |
| | | | first step, C | p429 | CATCCAAATGAAGATGTACAAACTGGTATCGTTCTG | delLinker2-3 (fw B) |
| | | | | | GTTCATTTCCGTATCTTGTTCTTG | Rev Nab2 C-term |
| | | | second step | N+C (from first step) | gtccactgGAATTCATGTCTCAAGAACAGTACACAGAAAA | Fwd for Nab2 ORF amplification with EcoRI site |
| | | | | | ACAAACGCGGCCGCGTTCATTTCCGTATCTTGTTCTTG | Rev NotI-[C-term of Nab2], no term codon, can be cloned in frame with FLAG in pESC |
| ΔL4-5 | p448 | [391-408], C-terminal flag-tag | first step, N | p429 | ATGTCTCAAGAACAGTACACAG | Forw Nab2 N-term |
| | | | | | AGGACTTTTCCGACAATGAAGAGTGGGC | delLinker4-5 (rev A) |
| | | | first step, C | p429 | CTTCATTGTCGGAAAAGTCCTTAGAACAATGTAAG | delLinker4-5 (fw B) |
| | | | | | GTTCATTTCCGTATCTTGTTCTTG | Rev Nab2 C-term |
| | | | second step | N+C (from first step) | gtccactgGAATTCATGTCTCAAGAACAGTACACAGAAAA | Fwd for Nab2 ORF amplification with EcoRI site |
| | | | | | ACAAACGCGGCCGCGTTCATTTCCGTATCTTGTTCTTG | Rev NotI-[C-term of Nab2], no term codon, can be cloned in frame with FLAG in pESC |
| NAB2-flag (wild type) | p434 | C-terminal flag-tag | n.a. | p429 | gtccactgGAATTCATGTCTCAAGAACAGTACACAGAAAA | Fwd for Nab2 ORF amplification with EcoRI site |
| | | | | | ACAAACGCGGCCGCGTTCATTTCCGTATCTTGTTCTTG | Rev NotI-[C-term of Nab2], no term codon, can be cloned in frame with FLAG in pESC |
| H434D | p481 | H434D | n.a. | p429 | TTCACGGCACATAATATCAGAACGAGCATGTCT | H434D-NAB2_fw |
| | | | | | GGAGCAAACTGTACTAGAATTGA | H434X-NAB2_rev |
| H434D, N466D | p482 | H434D, N466D | n.a. | p481 | TCTGAATAGACAGTAAATATCCTTACAATTGACACC | N466D-NAB2_fw |
| | | | | | CATCCTCCAGGCAGAGTAC | 466-468-NAB2_rev |
| F450A | p495 | F450A | n.a. | p429 | CATTAATTGGATGGCCAGCTAAACAATCAATTC | nab2-F450A fw |
| | | | | | AAGATTGTAGATTTGGTGTCAATTG | nab2_F450A_rev |
| F353A | p517 | F353A | n.a. | p429 | GCTGGTGTTGGATGACCAGCTGGGCATGATGGATTGG A | F353A_fw |
| | | | | | AAATGAAGATGCGAAAGTCATTG | 353_rev |
| F353A, H434D, N466D | p518 | F353A, H434D, N466D | n.a. | p482 | GCTGGTGTTGGATGACCAGCTGGGCATGATGGATTGG A | F353A_fw |
| | | | | | AAATGAAGATGCGAAAGTCATTG | 353_rev |
| F353A, F450A | p508 | F353A, F450A | n.a. | p495 | GCTGGTGTTGGATGACCAGCTGGGCATGATGGATTGG A | F353A_fw |
| | | | | | AAATGAAGATGCGAAAGTCATTG | 353_rev |
| K341A, F342A | p516 | K341A, F342A | n.a. | p429 | GAACACAGAGCCCCAGCAGCACACAGAACGATACCAG TTTG | K341A,F342A_fw |
| | | | | | CAATCCATCATGCCCATTTG | 342_rev |
| K341A, F342A, F450A | p507 | K341A, F342A, F450A | n.a. | p495 | GAACACAGAGCCCCAGCAGCACACAGAACGATACCAG TTTG | K341A,F342A_fw |
| | | | | | CAATCCATCATGCCCATTTG | 342_rev |
| H420D | p478 | H420D | n.a. | p429 | ACGTTTATTGGTGCAATCCGTACCGAACTTACA | H420D-NAB2_fw |
| | | | | | TGCAAATATAGACATGCTCGTTC | H420D-NAB2_rev |
| H420D, H434D, N466D | p496 | H420D, H434D, N466D | n.a. | p482 | ACGTTTATTGGTGCAATCCGTACCGAACTTACA | H420D-NAB2_fw |
| | | | | | TGCAAATATAGACATGCTCGTTC | H420D-NAB2_rev |
| H434D, N466D, I467D, Y468D | p497 | H434D, N466D, I467D, Y468D | n.a. | p482 | ATCATCATCCTTACAATTGACACCAAATC | I467D,Y468D on N466D_fw |
| | | | | | TGTCTATTCAGACATCCTCCAG | 468-rev |
| ZnF1-7-CT >> Hsa ZnF1-5 | p458 | SceNab2 amino acids 1-255 fused to HsaZC3H14 amino acids 594-736 | first step, N | p429 | ATGTCTCAAGAACAGTACACAG | Forw Nab2 N-term |
| | | | | | TGGAGTGAAGTTCATATTACTCTC | NAB2-255 rev |
| | | | second step | N (from first step) + GeneArt DNA fragment | gtccactgGAATTCATGTCTCAAGAACAGTACACAGAAAA | Fwd for Nab2 ORF amplification with EcoRI site |
| | | | | | ACAAACGCGGCCGCTCATTCGGAGGTTTGTGGTC | ZC3H14 Cterm rev-NotI |
| **p(*CEN/LEU2*) parent plasmids** | | | | | | |
| empty | p513 | n.a. | n.a. | pAC1039 | cctcgaggtcgacggtatcgataagcttgCGAGACGTTTATATAGGG ATGTG | NAB2-promoter_fw |
| | | | | | gcggccgccccgggctgcaggaattcgTTCTGATGTACTTCCACTT CCT | NAB2-promoter_rev |
| | | | n.a. | pAC1039 | cgaattcctgcagcccggggcggccgcTTACTATTAAAATCACGGA ACGAAATTC | NAB2-terminator_fw |

| | | | | | | |
|---|---|---|---|---|---|---|
| | | | | | taaagggaacaaaagctggagctccTGATTGAAACCCAGTCTGTCC | NAB2_terminator_rev |
| empty | pMT101 | autoregulatory A26>A11 | n.a. | p513H | GTTTTTAAACAGTTCCTGATC | NAB2-UTR_fw |
| | | | | | TTTTTTTTTTTAAAATCTAAATATGTTGCTTG | NAB2-UTR_A11_rev |
| empty | pMT102 | autoregulatory A26>A16 | n.a. | p513H | GTTTTTAAACAGTTCCTGATC | NAB2-UTR_fw |
| | | | | | TTTTTTTTTTTTTTTTAAAATCTAAATATGTTGCTTG | NAB2-UTR_A16_rev |
| empty | pMT103 | autoregulatory A26>A33 | n.a. | p513H | GTTTTTAAACAGTTCCTGATC | NAB2-UTR_fw |
| | | | | | TTTTTTTTTTTTTTTTTTTTTTTTTTTTTTTTTAAAATCTAAATATGTTGCTTG | NAB2-UTR_A33_rev |
| empty | pMT104 | autoregulatory A26>A42 | n.a. | p513H | GTTTTTAAACAGTTCCTGATC | NAB2-UTR_fw |
| | | | | | TTTTTTTTTTTTTTTTTTTTTTTTTTTTTTTTTTTTTTTTTTAAAATCTAAATATGTTGCTTG | NAB2-UTR_A42_rev |
| ***E. coli*** **expression plasmids** | | | | | | |
| Nab2 | pEG005 | pET28b C-terminal 6xHis | n.a. | p429 | CTTTAAGAAGGAGATATACCATGTCTCAAGAACAGTACACAG | Nab2-Nterm_homol to pET28b_Gibson |
| | | | | | TGGTGCTCGAGTGCGGCCGCTCAATGATGATGATGATGATGGTTCATTTCCGTATCTTGTT | PRIM047_Nab2-Cterm-6His_homol to pET28b_Gibson |
| Nab2-FKBP12 | pEG025 | pET28b C-terminal FKBP12 (H. sapiens FKBP12 residues 1-107), C-terminal 6xHis | n.a. | p429 | CTTTAAGAAGGAGATATACCATGTCTCAAGAACAGTACACAG | Nab2-Nterm_homol to pET28b_Gibson |
| | | | | | GTTCATTTCCGTATCTTGTT | NAB2 ORF rev |
| | | | n.a. | genomic DNA of HHY212 (Euroscarf) | GATACGGAAATGAACGGAGTGCAGGTGGAAACCATCTC | NAB2-3end_FKBP12_fw |
| | | | | | CTCGAGTGCGGCCGCTCAATGATGATGATGATGATGGCCAGTTTCCAGTTTTAGAAGCTC | FKBP12-6His_rev_pET28 overhand |
| Nab2-FRB | pEG026 | pET28b C-terminal FRB (H. sapiens mTOR residues 2021-2113, T2098L), C-terminal 6xHis | n.a. | p429 | CTTTAAGAAGGAGATATACCATGTCTCAAGAACAGTACACAG | Nab2-Nterm_homol to pET28b_Gibson |
| | | | | | GTTCATTTCCGTATCTTGTT | NAB2 ORF rev |
| | | | n.a. | pFA6a-FRB-GFP-KanMX6 (Euroscarf P30580) | AACAAGATACGGAAATGAACATCCTCTGGCATGAGATG | NAB2_3end_overhang_FRB-fw |
| | | | | | CTCGAGTGCGGCCGCTCAATGATGATGATGATGATGGCCCTTTGAGATTCGTCGGA | FRB-G-6his-stop-pET28b_rev |
| Nab2_ΔRGG | pEG007 | pET28b [201-255], C-terminal 6xHis | n.a. | p442 | CTTTAAGAAGGAGATATACCATGTCTCAAGAACAGTACACAG | Nab2-Nterm_homol to pET28b_Gibson |
| | | | | | TGGTGCTCGAGTGCGGCCGCTCAATGATGATGATGATGATGGTTCATTTCCGTATCTTGTT | PRIM047_Nab2-Cterm-6His_homol to pET28b_Gibson |
| Nab2_ΔZnF1-4 | pEG006 | pET28b [262-389], C-terminal 6xHis | n.a. | p440 | CTTTAAGAAGGAGATATACCATGTCTCAAGAACAGTACACAG | Nab2-Nterm_homol to pET28b_Gibson |
| | | | | | TGGTGCTCGAGTGCGGCCGCTCAATGATGATGATGATGATGGTTCATTTCCGTATCTTGTT | PRIM047_Nab2-Cterm-6His_homol to pET28b_Gibson |
| Nab2_ΔZnF5-7 | pEG012 | pET28b [390-485], C-terminal 6xHis | n.a. | pEG005 | GCACCCAATTCAAACGTT | deltaZnF567_fw |
| | | | | | CAATGAAGAGTGGGCCTT | deltaZnF567_rev |
| Nab2_N466D,H434D | pEG008 | pET28b N466D,H434D, C-terminal 6xHis | n.a. | p482 | CTTTAAGAAGGAGATATACCATGTCTCAAGAACAGTACACAG | Nab2-Nterm_homol to pET28b_Gibson |
| | | | | | TGGTGCTCGAGTGCGGCCGCTCAATGATGATGATGATGATGGTTCATTTCCGTATCTTGTT | PRIM047_Nab2-Cterm-6His_homol to pET28b_Gibson |
| Nab2_F450A | pEG009 | pET28b F450A, C-terminal 6xHis | n.a. | p495 | CTTTAAGAAGGAGATATACCATGTCTCAAGAACAGTACACAG | Nab2-Nterm_homol to pET28b_Gibson |
| | | | | | TGGTGCTCGAGTGCGGCCGCTCAATGATGATGATGATGATGGTTCATTTCCGTATCTTGTT | PRIM047_Nab2-Cterm-6His_homol to pET28b_Gibson |
| Nab2_F353A | pEG010 | pET28b F353A, C-terminal 6xHis | n.a. | p517 | CTTTAAGAAGGAGATATACCATGTCTCAAGAACAGTACACAG | Nab2-Nterm_homol to pET28b_Gibson |
| | | | | | TGGTGCTCGAGTGCGGCCGCTCAATGATGATGATGATGATGGTTCATTTCCGTATCTTGTT | PRIM047_Nab2-Cterm-6His_homol to pET28b_Gibson |
| Nab2_F353A,H434D,N466D | pEG017 | pET28b F353A,H434D,N466D, C-terminal 6xHis | n.a. | p518 | CTTTAAGAAGGAGATATACCATGTCTCAAGAACAGTACACAG | Nab2-Nterm_homol to pET28b_Gibson |
| | | | | | TGGTGCTCGAGTGCGGCCGCTCAATGATGATGATGATGATGGTTCATTTCCGTATCTTGTT | PRIM047_Nab2-Cterm-6His_homol to pET28b_Gibson |
| Nab2_F353A,F450A | pEG011 | pET28b F353A,F450A, C-terminal 6xHis | n.a. | p508 | CTTTAAGAAGGAGATATACCATGTCTCAAGAACAGTACACAG | Nab2-Nterm_homol to pET28b_Gibson |
| | | | | | TGGTGCTCGAGTGCGGCCGCTCAATGATGATGATGATGATGGTTCATTTCCGTATCTTGTT | PRIM047_Nab2-Cterm-6His_homol to pET28b_Gibson |

### Supplemental Table S4. Oligonucleotides

| Identifier | Name of RNA oligonucleotide | Sequence 5'-3' (underlined are deoxyribonucleotides) | 5' mod | 3' mod | Sequence reference | Manufactured by |
|---|---|---|---|---|---|---|
| R001 | *CYC142* | UUUAUAGUUAUGUUAGUAUUAAGAACGUUA UUUAUAUUUCAA | Atto680 | - | Schmid *et al.* 2012 | Eurogentec |
| R022 | *CYC159* | UUUAUAGUUAUGUUAGUAUUAAGAACGUUA UUUAUAUUUCAAAUUUUUCUUUUUUUUCU | Atto680 | Cy3 | *CYC1a* (Hill *et al.* 2019) extended by 3 nts at the 3'end | Splinted ligation of R001 and R021 with D004; 1:1.2:1; T4 DNA Ligase 0.4 U/μL |
| R021 | *dwCYC117* | AUUUUUCUUUUUUUUCU | Phosphate | Cy3 | This study | Eurogentec |
| R003 | *A59* | AAAAAAAAAAAAAAAAAAAAAAAAAAAAAAAA AAAAAAAAAAAAAAAAAAAAAAAAAAA | Atto680 | - | This study | Eurogentec |
| R006 | *A59* | AAAAAAAAAAAAAAAAAAAAAAAAAAAAAAAA AAAAAAAAAAAAAAAAAAAAAAAAAAA | Phosphate | - | This study | Eurogentec |
| R002 | *A30* | AAAAAAAAAAAAAAAAAAAAAAAAAAAAAA | Phosphate | - | This study | Eurogentec |
| R007 | *A15* | AAAAAAAAAAAAAAA | Atto680 | - | This study | Eurogentec |
| R017 | *A20* | AAAAAAAAAAAAAAAAAAAA | Atto680 | - | This study | Splinted ligation of R007 and R008 with D002, 1:1.5:1.5, T4 DNA Ligase 0.4 U/μL |
| R012 | *A25* | AAAAAAAAAAAAAAAAAAAAAAAAA | Atto680 | - | This study | Splinted ligation of R007 and R009 + R010 with D002, 1:1.5:1, T4 DNA Ligase 0.4 U/μL |
| R013 | *A30* | AAAAAAAAAAAAAAAAAAAAAAAAAAAAAA | Atto680 | - | This study | Splinted ligation of R007 and R009 + R010 with D002, 1:1.5:1, T4 DNA Ligase 0.4 U/μL |
| R014 | *A35* | AAAAAAAAAAAAAAAAAAAAAAAAAAAAAAAA AAA | Atto680 | - | This study | Splinted ligation of R007 and R009 + R010 with D002, 1:1.5:1, T4 DNA Ligase 0.4 U/μL |
| R016 | *A45* | AAAAAAAAAAAAAAAAAAAAAAAAAAAAAAAA AAAAAAAAAAAAA | Atto680 | - | This study | Splinted ligation of R007 and R009 + R010 with D002, 1:1.5:1, T4 DNA Ligase 0.4 U/μL |
| R008 | *A5* | AAAAA | Phosphate | - | This study | Eurogentec |
| R009 | *A10* | AAAAAAAAAA | Phosphate | - | This study | Eurogentec |
| R010 | *A15* | AAAAAAAAAAAAAAA | Phosphate | - | This study | Eurogentec |
| D002 | splint DNA | TTTTTTTTTTTTTTTTTTTTTTTTTTTTTTTTTTT TTTTTTTTTTTTTTTGAAATATAAATAA | - | - | This study | Merck/SigmaAldrich |
| D004 | splint DNA | AAAAGAAAAATTTGAAATATAAA | - | - | This study | Merck/SigmaAldrich |
| DL163 | *HSP104* RNaseH targeting oligo | ACATTTTCATCACGAGATTTACCC | - | - | Libri *et al.* 2002 | Merck/SigmaAldrich |
| DL164 | *HSP104* 3´hybridization DNA probe | TTATCGTCATCACCTAACGTGTCAGCCCCTA TAGTAGCTTCGTGATTTGGTAGAACTTCC | - | - | Libri *et al.* 2002 | Merck/SigmaAldrich |
| MS618 | *SCR1* hybridization DNA probe | GGTCACCTTTGCTGACGCTGG | - | - | Schmid *et al.* 2015 | Merck/SigmaAldrich |
| NL-A48 | nanolever green channel (switchSENSE) | TAGTGCTGTAGGAGAATATACGGGCTGCTCG TGTTGACAAGTACTGAT | - | green dye | Dynamic Biosensors | n.a. |
| cNLA-X2 | nanolever green channel (switchSENSE) | ACAGATGACTGCTCGTCAATATCAGTACTTG TCAACACGAGCAGCCCGTATATTCTCCTACA GCACTA | - | - | This study | Eurogentec |
| CX2 | DNA only (switchSENSE) | ATTGACGAGCAGTCATCTGT | - | - | This study | Eurogentec |
| NL-B48 | nanolever red channel (switchSENSE) | TAGTCGTAAGCTGATATGGCTGATTAGTCGG AAGCATCGAACGCTGAT | - | red dye | Dynamic Biosensors | n.a. |
| cNLB-X1 | nanolever red channel (switchSENSE) | ACACTACTGACGAGCACAATATCAGCGTTCG ATGCTTCCGACTAATCAGCCATATCAGCTTA CGACTA | - | - | This study | Eurogentec |
| CX1-rA60 | rA60 (switchSENSE) | ATTGTGCTCGTCAGTAGTGTAAAAAAAAAAAA AAAAAAAAAAAAAAAAAAAAAAAAAAAAAAAA AAAAAAAAAAAAAAAA | - | - | This study | Eurogentec |
| CX1-rA30 | rA30 (switchSENSE) | ATTGTGCTCGTCAGTAGTGTAAAAAAAAAAAA AAAAAAAAAAAAAAAAAA | - | - | This study | Eurogentec |

#### Supplemental Table S5. Polyadenylation reaction conditions.

| Figure panel | Pre-incubation step 1 | Pre-incubation step 2 | Final concentrations | | | | | |
|---|---|---|---|---|---|---|---|---|
| | | | RNA | CPF | CF IA | CF IB | Nab2-6xHis | ATP |
| 1B & 1D | RNA +CF IA +CF IB (5 min) | +CPF (3 min) | 50 nM | 50 nM | 225 nM | 225 nM | - | 2 mM |
| S1A | RNA + CPF (5 min) | - | 100, 50, 25 nM | 100, 50, 25 nM | 500 nM | 500 nM | - | 2 mM |
| S1B | RNA +CF IA +CF IB (5 min) | - | 25 nM | 75 nM | 500 nM | 500 nM | - | 2 mM |
| S1C | RNA +CF IA +CF IB (5 min) | - | 25 nM | 75 nM | 500 nM | 500 nM | - | 100 µM (3'dATP) |
| 2A | RNA +CF IA +CF IB (5 min) | +CPF -/+Nab2 (3 min) | 25 nM | 75 nM | 500 nM | 500 nM | 0, 500 nM | 2 mM |
| 2B | RNA +CF IA +CF IB (5 min) | +CPF -/+Nab2 (4 min) | 25 nM | 75 nM | 500 nM | 500 nM | 0, 50, 100, 175, 250, 350, 500, 750, 1000 nM | 2 mM |
| 2C | RNA +CF IA +CF IB (5 min) | +CPF -/+Nab2 (4 min) | 25 nM | 75 nM | 500 nM | 500 nM | 175, 350 nM | 500, 100, 50, 20, 10, 5 µM |
| 4C&S4B | RNA +CF IA +CF IB (6 min) | +CPF -/+Nab2 -/+ Rapamycin or DMSO (5 min) | 25 nM | 75 nM | 500 nM | 500 nM | 0, 175, 500, 1000 nM | 2 mM |
| 5C | RNA +CF IA +CF IB (5 min) | +CPF -/+Nab2 (3 min) | 25 nM | 75 nM | 500 nM | 500 nM | 0, 175, 250, 350, 500, 750, 1000 nM | 2 mM |
| 6B | RNA +CF IA +CF IB (5 min) | +CPF -/+Nab2 (3 min) | 25 nM | 75 nM | 500 nM | 500 nM | 0, 175, 250, 350, 500, 750, 1000 nM | 2 mM |
| S6B | RNA +CF IA +CF IB (5 min) | +CPF -/+Nab2 (3 min) | 25 nM | 75 nM | 500 nM | 500 nM | 0,1000 nM | 2 mM |
